# Infection-driven activation of transglutaminase 2 boosts glucose uptake and hexosamine biosynthesis

**DOI:** 10.1101/634501

**Authors:** Benoit Maffei, Marc Laverrière, Yongzheng Wu, Sébastien Triboulet, Stéphanie Perrinet, Magalie Duchateau, Mariette Matondo, Robert L. Hollis, Charlie Gourley, Jan Rupp, Jeffrey W. Keillor, Agathe Subtil

**Affiliations:** Institut Pasteur, Unité de Biologie cellulaire de l’infection microbienne, 25 rue du Dr Roux, 75015 Paris, France; CNRS UMR3691, Paris, France; Sorbonne Université, Collège Doctoral, F-75005 Paris, France; Plateforme Protéomique, Unité de Spectrométrie de Masse pour la Biologie, USR 2000 CNRS, Institut Pasteur, Paris, France; Nicola Murray Centre for Ovarian Cancer Research, Cancer Research UK Edinburgh Centre, MRC IGMM, University of Edinburgh, Edinburgh, UK; Department of Infectious Diseases and Microbiology, University of Lübeck, Lübeck, Germany; Department of Chemistry and Biomolecular Sciences, University of Ottawa, Canada

**Keywords:** Chlamydia, GFPT, Hexosamine biosynthesis, *O*-GlcNAcylation, Transglutaminase 2

## Abstract

Transglutaminase 2 (TG2) is a ubiquitous enzyme with transamidating activity. We report here that the expression and activity of TG2 are enhanced in cells infected with the obligate intracellular bacteria *Chlamydia trachomatis*. Genetic or pharmacological inhibition of TG2 activity impair bacterial development. We show that TG2 increases glucose import by up-regulating the transcription of the glucose transporter genes *GLUT-1* and *GLUT-3*. Furthermore, TG2 activation drives one specific glucose-dependent pathway in the host, i.e. hexosamine biosynthesis. Mechanistically, we identify the glucosamine:fructose-6-phosphate amidotransferase (GFPT) among the substrates of TG2. GFPT modification by TG2 increases its enzymatic activity, resulting in higher levels of UDP-N-acetylglucosamine biosynthesis. As a consequence, TG2 activation results in increased protein *O*-GlcNAcylation. The correlation between TG2 transamidating activity and *O*-GlcNAcylation is disrupted in infected cells because host hexosamine biosynthesis is being exploited by the bacteria, in particular to assist their division. In conclusion, our work establishes TG2 as a key player in controlling glucose-derived metabolic pathways in mammalian cells, themselves hijacked by *C. trachomatis* to sustain their own metabolic needs.

## INTRODUCTION

The enzyme transglutaminase 2 (TG2) is an extremely versatile protein exhibiting transamidase, protein disulfide isomerase and guanine and adenine nucleotide binding and hydrolyzing activities (Gundemir, Colak et al., 2012). Also designated as “tissue transglutaminase”, it is ubiquitously expressed in the cytoplasm and at the cell surface in association with the extracellular matrix (Eckert, Kaartinen et al., 2014). The transamidase activity is the best described activity, and it is regulated by Ca^2+^ (Gundemir et al., 2012). It results in the formation of cross-links between proteins, or of post-translational modification of a protein substrate through incorporation of a small primary amine, or deamidation of a glutamine into a glutamate. Under steady-state conditions, TG2 exists in a compact, inactive conformation. Increase in intracellular Ca^2+^ concentration (upon stress, cell activation, etc) causes a conformational change, and the enzyme becomes catalytically active as a transamidase. Studies of genetically engineered mouse models and/or inherited disorders have implicated TG2 in several pathological conditions (Iismaa, Mearns et al., 2009). In particular, increased TG2 expression and transamidation activity is a common feature of many inflammatory diseases and events (Eckert et al., 2014). Possibly linked to its increased expression in inflammatory processes, several lines of evidence suggest the involvement of TG2 during cancer development (Huang, Xu et al., 2015).

Surprisingly, while the association between TG2 activity and inflammatory situations has been studied in several normal and pathological situations (Di Sabatino, Vanoli et al., 2012, Huang et al., 2015, Ientile, Curro et al., 2015, Iismaa et al., 2009, Liu, Kellems et al., 2017), the implication of this enzyme in a very classical inflammatory process, e.g. the defense response of a tissue to the invasion by a microorganism, has remained very poorly investigated. *Chlamydia trachomatis* is the most common sexually transmitted bacterial pathogen, and it develops inside a vacuole in a human host cell, typically an epithelial cell of the genital tract (reviewed in (AbdelRahman & Belland, 2005)). This obligate intracellular bacterium depends on the host to supply several essential metabolites, and in particular glucose (Gehre, Gorgette et al., 2016, Stephens, Kalman et al., 1998). Epithelial cells respond to the infection with the secretion of proinflammatory cytokines such as IL-6 and IL-8 (Rasmussen, Eckmann et al., 1997). The inflammatory response is exacerbated upon reinfection, ultimately leading to tissue damage such as hydrosalpinx and fibrosis (Brunham & Rey-Ladino, 2005). In this work, we show that TG2 becomes activated during the infection of epithelial cells with *C. trachomatis*, and is required for optimal bacterial growth. The investigation of the consequence of TG2 activation on host metabolism and the identification of targets of TG2 transamidase activity during infection uncovered the control exerted by this enzyme on glucose import and on the hexosamine biosynthesis pathway, two metabolic features that are exploited by *C. trachomatis*.

## RESULTS

### TG2 is highly expressed and becomes active during *C. trachomatis* infection

A widely-used technique to probe TG2 activation is to measure the incorporation of biotin pentylamine (BP) into proteins. When present in excess, this membrane permeable primary amine out-competes other substrates for the transamidase reaction catalyzed by TG2 and becomes covalently linked to glutamine residues of TG2 substrate proteins. The biotin group is then easily detectable by western blot using streptavidin coupled to horseradish peroxidase (HRP) (Lee, Maxwell et al., 1992). This procedure was applied to HeLa cells infected or not for 48 h with *C. trachomatis*. In non-infected samples, BP incorporation was extremely low, as expected since in resting cells low Ca^2+^ concentration maintains TG2 in an inactive conformation (Gundemir et al., 2012). In contrast, infected cells showed a significant incorporation of BP. CP4d, an inhibitor of TG2 transamidating activity (Caron, Munsie et al., 2012), abolished BP incorporation in a dose-dependent manner, indicating that BP incorporation was the result of to the transamidase activity of TG2 (Fig. 1A). Live proliferating bacteria were needed for TG2 activation since filtered or heat-inactivated bacteria, or bacteria treated with the antibiotic doxycycline immediately after infection to prevent their proliferation, failed to induce BP incorporation (Fig. 1B and Fig. S1). BP incorporation upon infection was also observed in wild type mouse embryonic fibroblasts (MEFs) but not in MEFs isolated from *tgm2* knocked-out animals (TG2^−/−^), further supporting the implication of TG2 in this process (Fig. 1C).

**Figure 1.**
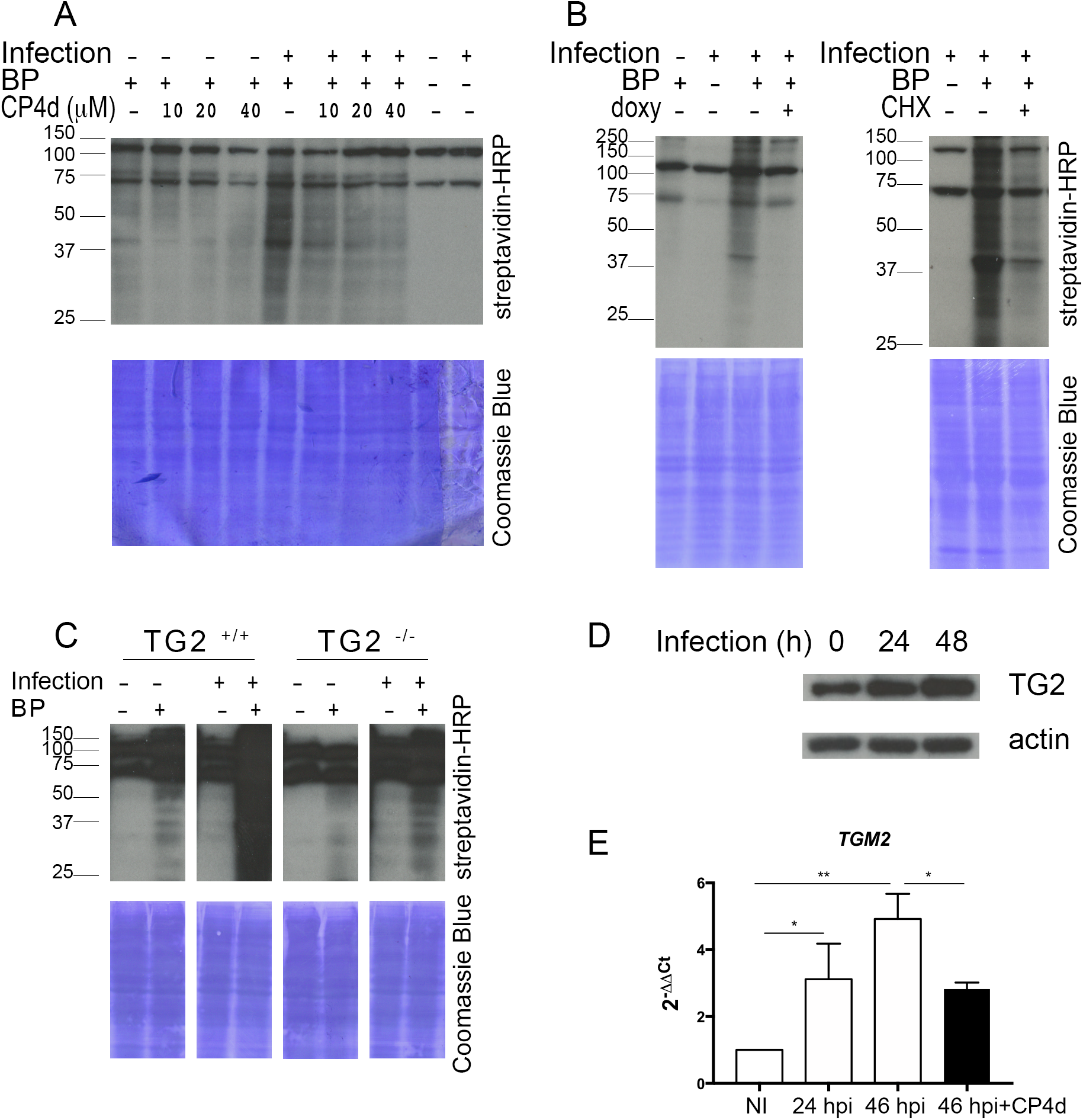
TG2 transamidase activity increases during *C. trachomatis* infection along with its expression. A – Whole cell lysates were prepared with HeLa cells infected or not for 48 h with *C. trachomatis* L2 (multiplicity of infection MOI=1) in the presence or not of BP. In the indicated samples CP4d was added 2 h before infection. Cell lysates were run on SDS-PAGE, proteins were transferred to a membrane and BP incorporation was revealed with HRP-conjugated streptavidin. BP incorporation is enhanced in infected samples, and is inhibited by CP4d. The two main bands present in all samples correspond to naturally biotinylated host proteins. After blotting the membrane was stained with Coomassie blue to control for equal loading. B – Same as in A, except that where indicated 250 μM doxycycline (doxy, left) or 7 μM cycloheximide (CHX, right) were added 24 h or 2 hpi, respectively. C – Whole cell lysates were prepared with TG2^+/+^ and TG2^−/−^ MEFs infected or not for 48 h with *C. trachomatis* L2 in the presence or not of BP, and analyzed as in A. D –Western blot with anti-TG2 antibodies on total cell lysates infected or not with *C. trachomatis* L2. E – Cells were infected with *C. trachomatis* L2 (MOI=1) for 24 or 48 h. Where indicated, 40 μM CP4d was added 2 hpi. *tgm2* transcripts were measured by real-time RT-PCR and normalized to *actin* transcripts following the ΔΔCt method. The data are presented as relative mRNA levels compared to uninfected cells and shown as the mean ± SD. Each experiment was performed in duplicate and repeated at four times. The asterisk indicates significantly different values (P < 0.05, Student’s ratio-paired t-test).

Probing cellular lysates using anti-TG2 antibodies showed that activation of TG2 was accompanied with an increased expression of the enzyme (Fig. 1D). Consistent with this observation, inhibition of protein synthesis with cycloheximide decreased infection-induced BP incorporation (Fig. 1B). Reverse transcription followed by quantitative PCR (RT-qPCR) measurements revealed a 3- to 4-fold increase in TG2 transcripts in infected versus non-infected cells, demonstrating that the increase in TG2 amount during infection is at least partly controlled at the transcriptional level (Fig. 1E). Interestingly, a positive feedback loop controls TG2 expression since TG2 transcription was no longer enhanced by infection when cells were treated with CP4d (Fig. 1E).

In conclusion, *C. trachomatis* infection increases TG2 levels and activates its transamidase activity.

### TG2 activity sustains bacterial growth

To determine if TG2 activity affected bacterial development, we infected HeLa cells in the presence or not of the transamidase inhibitor CP4d. Thirty h later, the progeny was collected and the number of infectious bacteria was determined by infecting fresh cells. Inhibition of TG2 activity with 40 μM CP4d resulted in a 2-fold decrease in bacterial progeny (Fig. 2A). Consistently, a similar reduction in bacterial progeny was observed when bacteria were grown on TG2^−/−^ MEFs compared to TG2^+/+^(Fig. S2A). To confirm these findings in primary cells, we used epithelial cells isolated from fallopian tubes (Roth, Konig et al., 2010). CP4d was even more potent at reducing the progeny after one developmental cycle in these cells than in HeLa cells, as a ten-fold reduction was observed for 10 μM CP4d (Fig. 2B). The negative impact of TG2 inhibition on bacterial development was also observed in primary cells infected with *C. trachomatis* serovar D, showing that the effect is not restricted to the LGV biovar (Fig. 2B).

**Figure 2.**
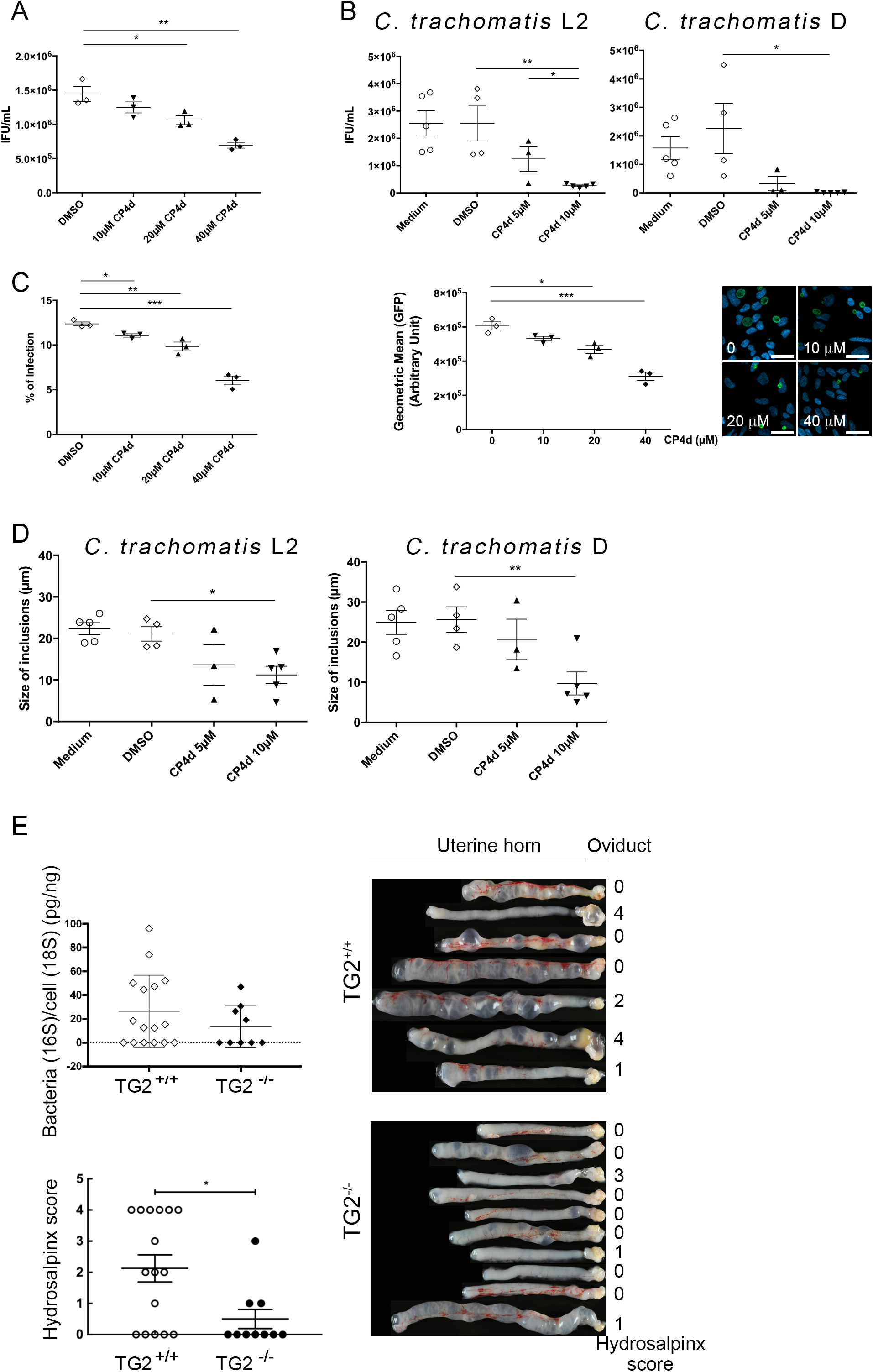
TG2 activity is needed for optimal *C. trachomatis* developmental and enhances hydrosalpinx upon *C. muridarum* infection in mice. A – HeLa cells were pre-treated with the indicated concentrations of CP4d (or DMSO alone) for 2 h before being infected with L2^incD^GFP at MOI=0.15. Thirty hours later the cells were disrupted and bacterial titers (IFU=inclusion forming unit) were determined by re-infecting fresh HeLa cells as described in the methods. The mean ± SD of three independent experiments are shown. B – Same as in A except that *C. trachomatis* serovar L2 (left) or D (right) were grown in primary cells isolated from fallopian tubes. For serovar D, IFU were determined 48 hpi. C – HeLa cells were pre-treated with the indicated concentrations of CP4d (or DMSO alone) for 2 h before being infected with L2^incD^GFP at MOI=0.15. Thirty hours later the cells were fixed and analyzed by flow cytometry. The percentage of infected cells (left) and the mean fluorescence of the infected population (right) are shown for three independent experiments. A representative field for each condition is shown, bar = 10 μM. D – Primary epithelial cells isolated from fallopian tubes were pre-treated with the indicated concentrations of CP4d (or DMSO alone) for 2 h before being infected with *C. trachomatis* serovar L2 (left) or D (right). Twenty-four hours later the cells were fixed, bacteria were stained using FITC-labeled anti-Chlamydia-LPS antibodies, and the size of the inclusions manually determined using ImageJ, on twenty inclusions per experiment. Asterisks indicate two significantly different values (*: P < 0.05; **: P < 0.01, Student’s paired t-test). E – Mice were infected intravaginally with 10^5^ IFU of *C. muridarum*. Twenty-five days later the mice were sacrificed and the upper genital tract, from the uterine horn to the oviduct, was collected. The right part was used for bacterial burden assessment (top left). The left part was rinsed with PBS and observed with a binocular magnifier (right) to determine the hydrosalpinx score (bottom left). Asterisk indicates significantly different values (P < 0.05, Student’s unpaired t-test).

Reduced progeny could result from impairment of one or several of the steps of the chlamydial developmental cycle: adhesion, entry, differentiation into the replicative form, proliferation, and differentiation into the infectious form. We observed that inhibition of TG2 had no effect on bacterial adhesion (Fig. S2B), but that 40 μM CP4d decreased the percentage of infected cells by about 2-fold in HeLa cells (Fig. 2C), consistent with a lower internalization efficiency (Fig. S2C). In addition, inclusions were smaller in cells treated with CP4d, and contained less bacteria, indicating that bacterial growth was slower in the absence of TG2 activity (Fig. 2C). The reduction of inclusion size when TG2 was inhibited was also observed in primary cells infected with *C. trachomatis* L2 or serovar D (Fig. 2D).

We next tested the incidence of the absence of TG2 on chlamydial development in a mouse model of infection. *Chlamydia muridarum* is a mouse-adapted strain genetically very close to *C. trachomatis* (Read, Brunham et al., 2000), and infection of HeLa cells with *C. muridarum* also activated TG2 (Fig. S3). We infected TG2^+/+^ and TG2^−/−^ mice intravaginally with *C. muridarum* and 25 days after infection, mice were sacrificed and the genital tract was isolated (Fig. 2E). DNA was extracted from the upper genital tract and bacterial load was determined by qPCR. A slightly higher number of mice retained detectable bacterial DNA in the wild type group (10/16, 62 %, for the TG2^+/+^ and 4/9, 44% for the TG2^−/−^ mice). Among the animals in which bacterial DNA was still detected, the trend was for a higher bacterial load in the wild type background, but the number of TG2^−/−^ animals we could breed was too low for statistical significance. These data indicate that the absence of TG2 reduces only marginally, if at all, the ability for *C. muridarum* to establish an infection. It is however possible that in some tissues the loss of TG2 is compensated by expression of other transgluatminases, limiting the interpretation of these data (Iismaa et al., 2009). One clear difference between the two groups came from anatomical observations: TG2^−/−^ animals showed milder signs of inflammation than their wild type littermate, especially when the oviduct hydrosalpinx scores were compared. However, it appears that the presence of TG2 exacerbates the tissue damage in this mouse model of infection, in line with the implication of TG2 in tissue fibrosis (Eckert et al., 2014, Iismaa et al., 2009).

### TG2 plays a central role in metabolic rewiring

We have recently shown that *C. trachomatis* acts as a glucose sink (Gehre et al., 2016). The host cell responds to glucose demand by increasing glucose uptake through overexpression of plasma membrane glucose transporters (Ojcius, Degani et al., 1998, Wang, Hybiske et al., 2017). Since we observed that TG2 level was increased during *C. trachomatis* infection, we wondered if this increase could control the concomitant increase in glucose transporter expression, as it does in mammary epithelial cells (Kumar, Donti et al., 2014). If this hypothesis was correct, one prediction that we could make was that low glucose availability in the culture medium should be more detrimental to bacterial growth in TG2^−/−^ MEFs compared to the wild type MEFs, as TG2^−/−^ MEFs would be impaired in their ability to adjust their glucose uptake to sustain bacterial growth. To test this hypothesis, we grew MEFs in medium containing decreasing concentrations of glucose and measured the number of infectious bacteria collected 30 hpi. As expected we observed that decreasing glucose availability resulted in a sharp decrease in bacterial titers, in both cellular backgrounds. However, bacterial titers were more sensitive to glucose deprivation in the TG2^−/−^ MEFs than in the wild-type cells (Fig. 3A). For instance, at 1 mg/mL glucose, the progeny was reduced by 82% in the TG2^−/−^ MEFs, compared to only 35% in the TG2^+/+^ MEFs.

**Figure 3.**
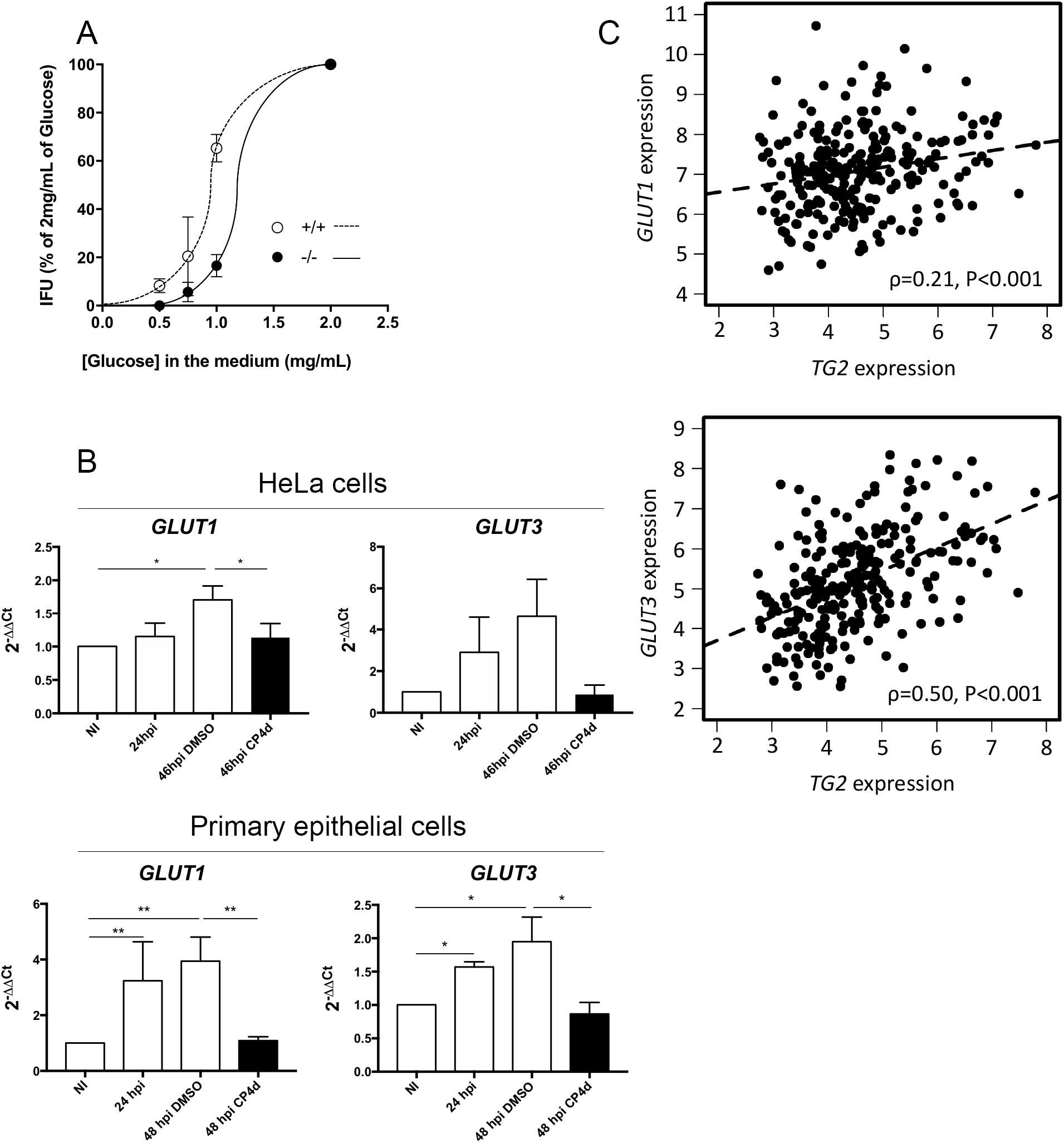
TG2 controls glucose import. A – MEFs were grown for 24 h culture medium complemented with the indicated concentration of glucose before being infected with L2 ^IncD^GFP bacteria (MOI = 0.2). Cells were disrupted 30 h later and the bacterial titer determined by re-infecting fresh wild type cells. The mean ± SD of three independent experiments are shown. B – Cells were infected with *C. trachomatis* L2 (MOI=1) for 24 or 48 h. Where indicated 40 μM CP4d was added 2 hpi. *GLUT-1* and *GLUT-3* transcripts were measured by real-time RT-PCR and normalized to *actin*. The data are presented as relative mRNA levels compared to uninfected cells and shown as the mean ± SD. Each experiment was performed in duplicate and repeated four times. Asterisks indicate two significantly different values (*: P < 0.05; **: P < 0.01, Student’s ratio-paired t-test). C — Relationship between *TGM2* and *GLUT-1* (top) and *GLUT-3* (bottom) expression across 265 HGSOCs. Expression comparisons were performed using Spearman’s rank correlation test.

To test the implication of TG2 in the rewiring of host metabolism more directly, we measured the incidence of TG2 inactivation on the cell capacity to uptake glucose. HeLa cells infected with *C. trachomatis* show increased transcription of *GLUT-1* and *GLUT-3*, which allows to increase glucose uptake and meet bacterial needs (Wang et al., 2017). We reproduced this result in HeLa cells, as well as in primary cells isolated from the endocervix (Fig. 3B). In contrast, in the presence of the TG2 inhibitor CP4d, the transcription of the glucose transporter genes was no longer induced by infection, indicating that TG2 is necessary for the control of *GLUT-1* and *GLUT-3* transcription. Absence of increase in *GLUT-1* and *GLUT-3* transcripts 48 hpi in the presence of CP4d was not due to the lower bacterial burden because when bacterial proliferation was interrupted 24 hpi by addition of doxycycline we observed a comparable reduction in bacterial load at 48 hpi as in cells treated with CP4d, but the transcription of the glucose transporter genes remained as high as in non-treated cells (Fig. S4).

Finally, to explore further the incidence of TG2 expression on that of glucose transporters we examined transcriptional data from a cohort of high grade serous ovarian cancer (HGSOC) patients. This population was chosen because clinical and biological data indicate that TG2 overexpression is an adverse prognostic factor in ovarian carcinoma (Hwang, Mangala et al., 2008, Shao, Cao et al., 2009). We observed a significant correlation between expression of *TGM2* and *GLUT-3* across the 265 clinical HGSOC specimens (ρ=0.50, P<0.001) (Fig.3C). The HGSOC cohort also demonstrated significant correlation between TG2 and GLUT-1 expression, though the magnitude of correlation was less marked (ρ=0.21, P<0.001) (Fig. 3C). Collectively, these data support the notion that TG2 plays a central role in regulating glucose transporters expression regulation in the context of infection or malignancy, thereby playing a central role in the control of the metabolic balance.

### TG2 targets glutamine:fructose-6-P amidotransferase and enhances its activity

In addition to its role in the up-regulation of the transcription of glucose transporter genes, TG2 may confer other benefits to *C. trachomatis*. To identify TG2 targets in the infectious process, HeLa cells were infected in the presence or absence of BP. Forty-eight hours later the cells were lysed and biotinylated proteins were isolated on streptavidin-coated beads, and identified by mass-spectrometry. Sixty-two proteins were found to be significantly enriched in the infected cell lysates grown in the presence of BP (Table S1). Fibronectin, galectin 3, RhoA, 40S ribosomal protein SA, immunoglobulin κ chain C region and hemoglobin beta were already identified as TG2 substrates (Guilluy, Rolli-Derkinderen et al., 2007, Mehul, Bawumia et al., 1995, Nelea, Nakano et al., 2008, Orrù, Caputo et al., 2003, Pincus & Waelsch, 1968, Sohn, Chae et al., 2010). BAG2 and several other mitochondrial proteins were also enriched in the samples prepared in the presence of BP, in agreement with TG2 being present and active in this compartment (Altuntas, Rossin et al., 2015).

Among the potential TG2 substrates we identified in *C. trachomatis* infected cells, the enzyme glutamine:fructose-6-P amidotransferase (GFPT) caught our attention because it uses fructose-6-P as a substrate, which is derived from glucose-6-P. Moreover, both isoforms of the enzyme, GFPT1 (also called GFAT) and GFPT2, had been recovered from the proteomic approach, making it a very strong hit. We first confirmed that GFPT was recovered in the biotinylated fraction of cells infected with *C. trachomatis* in the presence of BP using anti-GFPT antibodies. The abundance of GFPT in the biotinylated fraction strongly decreased when infection had been performed in the presence of the TG2 inhibitor CP4d, demonstrating that incorporation of the biotinylated probe in GFPT depended on the activity of TG2 (Fig. 4A).

**Figure 4.**
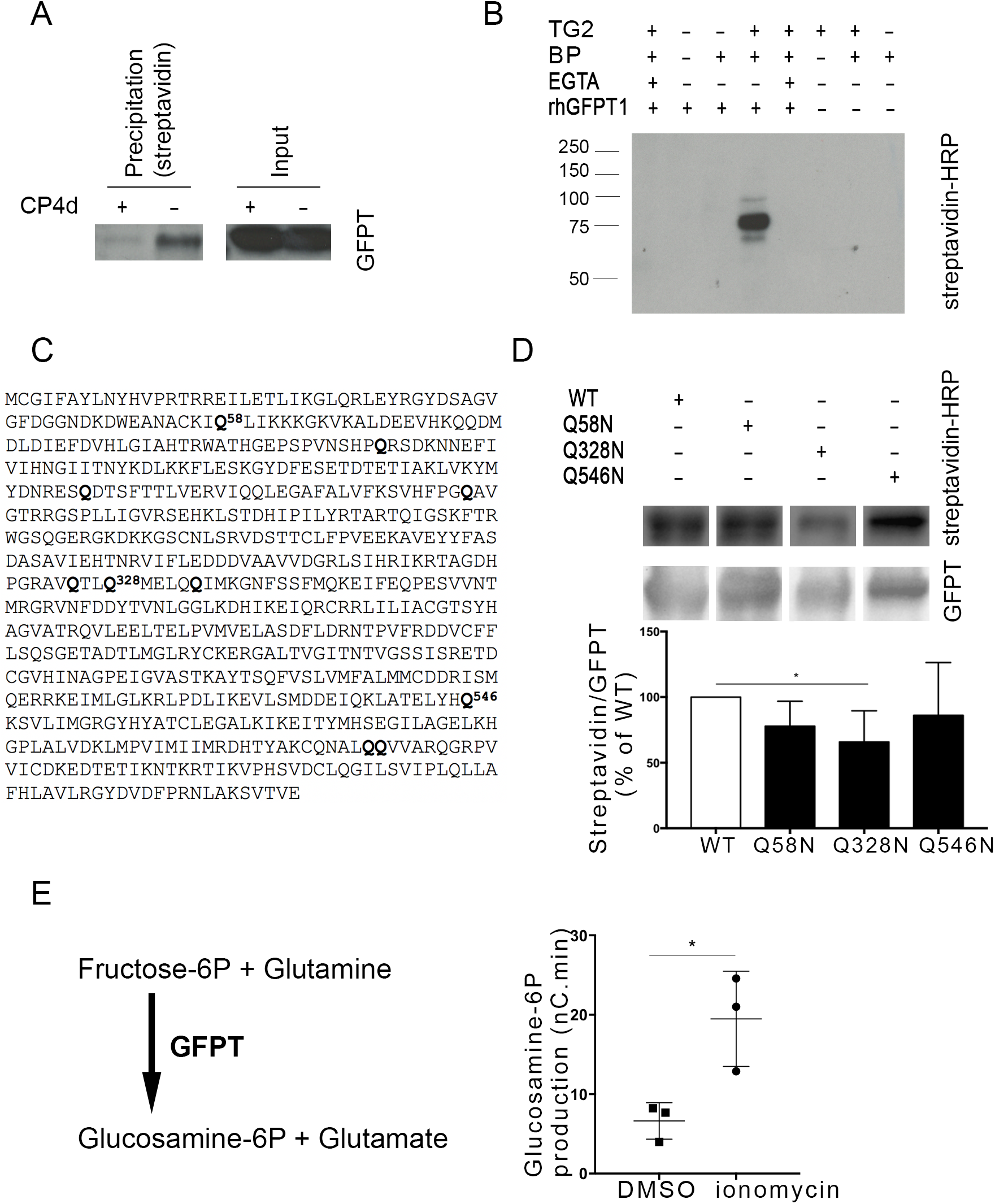
GFPT is a substrate of TG2 transamidase activity. A – HeLa cells were infected with *C. trachomatis* (MOI = 1) and 40 μM CP4d was added or not 2 hpi. After 24 h 0.5 mM BP was added and cells were lysed at 48 hpi. Lysates were precipitated with streptavidin-coated beads. After separation with SDS-PAGE, proteins were transferred to a membrane and blotted with anti-GFPT antibody followed with HRP-conjugated secondary antibody. B – *In vitro* assay testing the ability of purified TG2 to crosslink purified rhGFPT1 with BP. Samples were incubated for 3 h at 37°C before separation by SDS-PAGE. Proteins were transferred to a membrane and BP was revealed using HRP-conjugated streptavidin. rhGFPT1 is 77.5 kDa. C – GFPT1 sequence: glutamine residues identified by mass spectrometry as cross-linked to BP are in bold letter. D – *In vitro* assay was performed as described in B using wild type rhGFPT1 (WT), rhGFPT1 Q58N, rhGFPT1 Q328N or rhGFPT1 Q546N as substrates. The reaction was performed at 37 °C for 30 min. After probing with HRP-Streptavidin the membrane was washed and probed with anti-GFPT antibodies followed with HRP-conjugated secondary antibodies. The ratio of modified protein (streptavidin signal) to the total GFPT is shown, normalized to its value with WT rhGFPT1. The mean of five independent experiments is shown, the asterisk indicates two significantly different values (P < 0.05, Student’s ratio-paired t-test). E – Lysates of cells treated or not for 6 h with ionomycin were incubated at 37 °C for 45 min with fructose-6-P and glutamine. The production of glucosamine-6-P was measured using HPAEC-PAD. Results of three independent experiments are shown, with mean and SD. The asterisk indicates two significantly different values (P<0.05, Student’s paired t-test).

To further validate that GFPT is a novel substrate of TG2, purified TG2 and recombinant human GFPT1 (rhGFPT1) were incubated for 3 h at 37 °C in the presence of BP as primary amine donor. The incorporation of the biotinylated probe was analyzed by blotting with HRP-coupled streptavidin. The biotinylated probe was incorporated into rhGFPT1 in the presence and not in the absence of TG2. Furthermore, chelation of Ca^2+^ by EGTA inhibited the incorporation of the probe, as expected for a reaction dependent on the transamidase activity of TG2 (Fig. 4B). We concluded from these experiments that GFPT is a novel substrate of TG2 that becomes modified by the transamidase activity of the enzyme during *C. trachomatis* infection.

In order to determine which glutamine residue(s) of GFPT1 was modified by TG2 *in vitro*, we analyzed the products of the reaction by mass spectrometry. BP incorporation was identified in ten glutamine residues (out of twenty-eight, Fig. 5C), presumably because promiscuous reactions occur *in vitro*. Among those, two glutamine residues were identified as prone to modification by TG2 using bioinformatics tools designed to score the peptidic environment favorable for TG2 activity, namely Q328 and Q546 (Keresztessy, Csosz et al., 2006, Sugimura, Hosono et al., 2006). We thus generated a glutamine to asparagine point mutant for each of these residues to minimize the impact on protein folding. As a control, we also mutated Q58, another candidate target identified by mass spectrometry but not surrounded by a consensus sequence for TG2. Purified recombinant proteins were incubated with TG2 and BP for 30 min at 37 °C before stopping the reaction. BP incorporation was significantly reduced only in the rhGFPT1 Q328N, indicating that the Q328 is a prominent glutamine for modification by TG2 (Fig. 4D).

**Figure 5.**
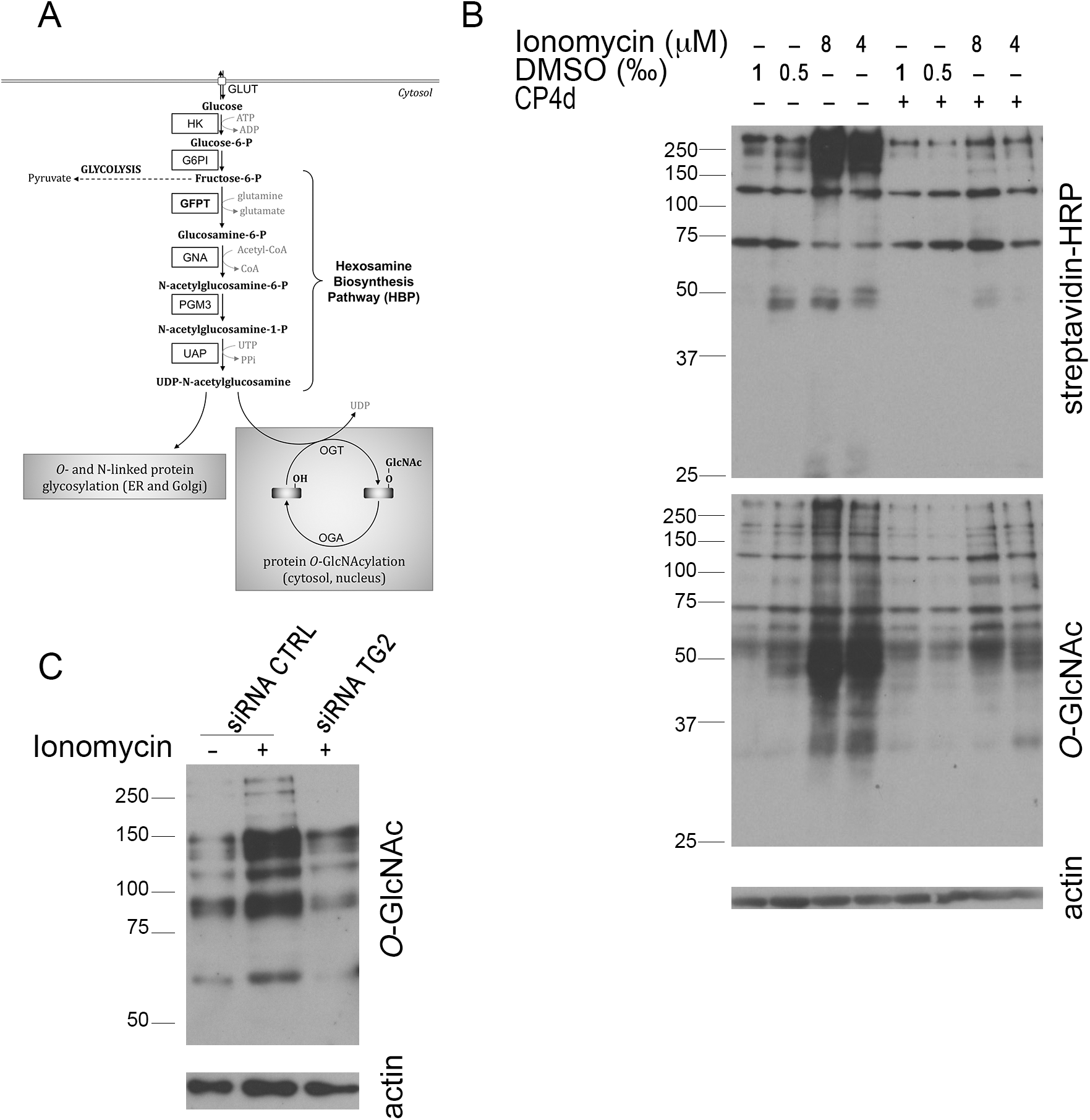
TG2 activation results in increased UDP-GlcNAc production. A – Schematic view of the hexosamine biosynthesis pathway. Production of glucosamine-6-P by GFPT is the first and rate-limiting step of the pathway that produces UDP-GlcNAc. HK: hexokinase; G6PI: glucose-6-P isomerase; GFPT: glutamine:fructose-6-P amidotransferase; GNA: glucosamine-6-P N-acetyltransferase; PGM3: phosphoglucomutase 3; UAP: UDP-N-acetylglucosamine pyrophosphorylase; OGT: *O*-GlcNAc transferase OGA: *O*-GlcNAcase; GlcNAc: N-acetylglucosamine. B – Endocervical epithelial cells were pre-treated or not with 40 μM CP4d for 2 h before addition of the indicated concentration of ionomycin (or an equivalent volume of DMSO) and 0.5 μM BP. Six hours later whole cell lysates were analyzed by western blot. The membrane was first blotted with HRP-conjugated streptavidin to detect TG2 activity, then extensively washed and probed with anti-*O*-GlcNAcylation antibody followed with HRP-conjugated secondary antibodies. Last the membrane was probed with anti-actin as a loading control. C – The same experimental procedure as described in B- was applied to HeLa cells treated for 48 h prior to ionomycin treatment (8μM) with siRNA control (siRNA CTRL) or siRNA directed against TG2 (siRNA TG2).

The fact that *C. trachomatis* produce their own GFPT prevented us from measuring the consequence of TG2 activation on host GFPT activity in infected cells. However, ionomycin is a widely used TG2 activator, as this Ca^2+^ ionophore increases intracellular Ca^2+^ concentration, which opens TG2 in its active conformation. We thus measured GFPT activity in lysates of cells treated or not with ionomycin, and analyzed the reaction products by high performance anion exchange chromatography (Fig. S5). We observed a three-fold increase in GFPT activity in cells treated with ionomycin, indicating that GFPT modification by TG2 increases the activity of the enzyme (Fig. 4E).

### Modification of GFPT by TG2 enhances the hexosamine biosynthesis pathway

The reaction catalyzed by GFPT is the first and rate limiting step of the hexosamine biosynthesis pathway (HBP, Fig. 5A). The HBP leads to the formation of uridine 5’-diphospho-*N*-acetylglucosamine (UDP-GlcNAc), which is further used for *N*-glycosylation, *N*-glycan branching, and *O*-linked *N*-acetylglycosylation (*O*-GlcNAcylation) in the ER, Golgi, and nucleus/cytosol, respectively. *O*-GlcNAcylation involves the transfer of a single UDP-GlcNAc moiety to the hydroxyl groups of serine or threonine residues. Two enzymes, *O*-GlcNActransferase (OGT) and *O*-GlcNAcase (OGA), catalyze *O*-GlcNAc addition and removal, respectively and the *O*-GlcNAc modification level of proteins is directly dependent on the concentration of UDP-GlcNAc, the donor substrate for OGT (Love & Hanover, 2005). To confirm that GFPT modification by TG2 increased its activity and thus hexosamine biosynthesis we measured the level of *O*-GlcNAcylation in primary epithelial cells. We observed an increase in *O*-GlcNAcylation in cells treated with ionomycin. This increase was dependent on TG2 activity since it was not observed in the presence of the TG2 inhibitor CP4d (Fig. 5B) or in cells in which *tgm2* expression had been silenced using siRNA (Fig. 5C). Altogether these experiments show that activation of TG2 transamidase activity enhances the HBP.

### The increase in the hexosamine biosynthetic pathway is hijacked by the bacteria

Surprisingly, we did not observe an increase in *O*-GlcNAcylation in cells infected for 48 h by *C. trachomatis* (Fig. 6A). Since *O*-GlcNAcylation directly depends on UDP-GlcNAc concentration this observation suggests that UDP-GlcNAc levels in the cytoplasm are not significantly increased in infected cells. We have previously demonstrated that *C. trachomatis* co-opts SLC35D2, a host antiporter transporting UDP-GlcNAc, UDP-glucose and GDP-mannose to import these metabolites into the vacuole in which the bacteria develop (Gehre et al., 2016). We reasoned that UDP-GlcNAc might not accumulate in the cytoplasm in infected cells because it was relocated to the inclusion lumen. Supporting this hypothesis, we observed that activation of TG2 by ionomycin elicited a lower increase in *O*-GlcNAcylation in infected cells compared to non-infected cells, indicating that less free UDP-GlcNAc is available for *O*-GlcNAcylation in the infected host cytoplasm (Fig. 6B). Importantly GFPT expression was stable in all conditions, indicating that the decrease in *O*-GlcNAcylation in infected cells is not due to lower GFPT expression (Fig. 6A-B).

**Figure 6.**
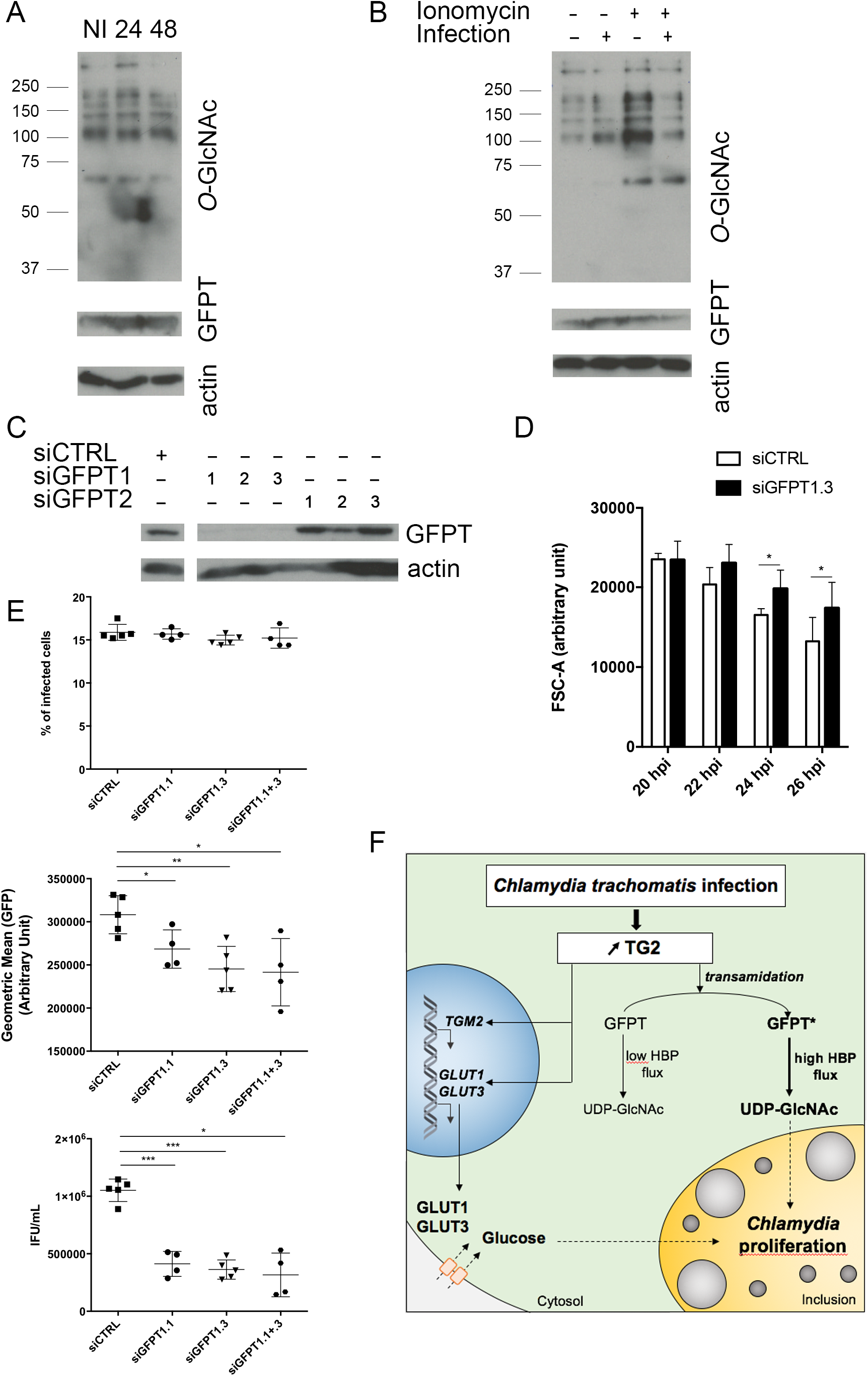
Optimal bacterial growth requires GFPT and prevents UDP-GlcNAc accumulation. A – HeLa cells were infected or not (NI) with *C. trachomatis* (MOI = 1), then lysed 24 or 48 hpi. After separation with SDS-PAGE, proteins were transferred to a membrane, probed with anti-*O*-GlcNAcylation antibody followed with HRP-conjugated secondary antibodies. After extensive washes the membrane was blotted again with anti-GFPT and anti-actin antibodies before revelation with HRP-conjugated secondary antibodies. B – HeLa cells were infected or not with *C. trachomatis* (MOI = 1). Twenty-four h later, 8 μM ionomycin (or DMSO alone) and 0.5 mM BP were added. After 6 h of treatment, cells were lysed and proteins revealed as in A. C – HeLa cells treated for 72 h with siRNA targeting GFPT1 or GFPT2 were lysed. After separation with SDS-PAGE, proteins were transferred to a membrane, probed with anti-GFPT and anti-actin antibody before revelation with HRP-conjugated secondary antibodies. D – HeLa cells were infected for the indicated times (MOI=0.3) before fixation, rupture of the cells and measurement of bacterial diameter. The mean diameter ± SD is shown, the asterisk indicates two significantly different values (P < 0.05, Student’s paired t-test). E – HeLa cells treated for 48 h with siRNA targeting GFPT1 or not (siCTRL) were infected with *C. trachomatis* (MOI = 0.15). Thirty hours later cells were fixed and analyzed by flow cytometry. The percentage of infected cells (top) and the mean fluorescence of the infected population (middle) are shown for at least four independent experiments. Duplicate wells were lysed and used to re-infect fresh HeLa cells to determine the bacterial titer (bottom). Asterisks indicate significantly different values (*: P < 0.05; **: P < 0.01; ***: P< 0.005, Student’s ratio-paired t-test). F – Schematic view of the outcome of TG2 activation in infection. The increase in TG2 expression and activity in cells infected with *C. trachomatis* results in the up-regulation of the expression of glucose transporters. Increasing quantities of glucose are thus imported in the host cytoplasm and redirected to the vacuole, where they fuel bacterial growth. Parallel to this transcriptional outcome, the transamidating activity of TG2 targets the host enzyme GFPT, thereby boosting the hexosamine biosynthesis pathway. The bacteria consume the resulting UDP-GlcNAc, or an intermediate along this pathway, in particular to sustain bacterial division.

In Gram-negative bacteria, UDP-GlcNAc supply is mostly used for lipopolysaccharide and peptidoglycan biosynthesis. *C. trachomatis* do not have a classical cell wall but use peptidoglycan synthesis for bacterial division (Liechti, Kuru et al., 2016). If UDP-GlcNAc, or an intermediate along the hexosamine biosynthesis pathway, was consumed at least partly in making bacterial peptidoglycan, lowering hexosamine biosynthesis should delay bacterial division, resulting in larger bacteria being formed. We tested this hypothesis by measuring the consequence of silencing GFPT, the rate-limiting enzyme in hexososamine biosynthesis, on bacterial size. The mass spectrometry data showed that both isoforms GFPT1 and GFPT2 were expressed in HeLa cells (Table S1). Comparison of the efficiency of siRNA designed to target specifically one isoform showed that GFPT2 was hardly detectable and targeting GFPT1 was sufficient to strongly decrease GFPT expression (Fig. 6C). We thus lowered hexosamine biosynthesis by treating cells with a siRNA against GFPT1 prior to infection, fixed the cells at increasing time of infection and used flow cytometry to measure bacterial sizes. As expected, the number of replicative bacteria increased with infection time (Fig. S5). Furthermore, it was recently demonstrated that replicative bacteria gradually decrease in size over the course of the developmental cycle (Lee, Enciso et al., 2017). We indeed observed a decrease in bacterial diameter over a 20 to 26 hpi time course, thereby validating the use of flow cytometry to measure the size of *C. trachomatis* (Fig. S6 and 6D). In cells treated with a siRNA against GFPT1 the mean bacterial diameter became significantly higher than in control cells 24 hpi (Fig. 6D). We confirmed this result by measuring the diameter of bacteria on electron microscopy pictures of cells infected for 30 h (Fig. S7). These kinetics fit well with our observation that TG2 activity increases between 24 and 48 hpi, when bacterial load is high, and access to nutrients might become limiting (Rother, Gonzalez et al., 2018). These observations support the hypothesis that a product of the hexosamine biosynthesis pathway is captured by the inclusion to support division. Consistent with a role for GFPT activity in sustaining bacterial growth we observed a reduction in the number of bacteria per inclusion 24 h post infection in the cells treated with siRNA against GFPT1, and the progeny collected was reduced 3-fold (Fig. 6E). Of note, silencing of GFPT1 had no incidence on bacterial entry and the initiation of bacterial development, as the percentage of infected cells was identical to that in control cells (Fig. 6E). Altogether, these data strongly support the hypothesis that the increase in hexosamine biosynthesis by the host upon GFPT modification by TG2 is exploited by the bacteria, in particular to assist bacterial division.

## DISCUSSION

TG2 transamidase activity is very potent, and would be deleterious if not tightly controlled. In basal conditions it is mostly turned off, and it is thought that under specific stress conditions, the enzyme might be locally turned on and transamidate specific substrates. The infection with *C. trachomatis* provided a unique physiological situation where the expression and activity of the enzyme were increased over a relatively short period of time in a physiological set-up. We took advantage of this observation to identify TG2 substrates. Among the 62 candidates identified, we focused on the enzyme GFPT. We showed that GFPT modification by TG2 increased its activity, resulting in higher hexosamine biosynthesis, a process also fueled by the positive control exerted by TG2 on glucose transporters expression. The product of the HBP, UDP-GlcNAc, is used for post-translational modification of proteins by *O*-GlcNAcylation. Thus, our work uncovered an unsuspected link between TG2 transamidase activity and *O*-GlcNAcylation. This link was disrupted in infected cells because the increase in hexosamine biosynthesis in the host was exploited by the bacteria, in particular to assist their division. In conclusion, our work establishes TG2 as a key player in controlling glucose-derived metabolic pathways in mammalian cells, themselves hijacked by *C. trachomatis* to sustain their own metabolic needs (Fig. 6F).

Several inflammatory conditions are associated with an increase in TG2 expression (Eckert et al., 2014). The promoter region of the TG2 gene (*tgm2*) contains, among others, response elements to NF-κB and to the pro-inflammatory cytokine interleukin-6 (IL-6), accounting for its increased transcription in inflammatory situations (Eckert et al., 2014, Suto, Ikura et al., 1993). The increase in TG2 transcription during *C. trachomatis* infection is likely due to the inflammatory response of epithelial cells (Rasmussen et al., 1997). How the enzyme becomes activated is less clear. In the cell Ca^2+^ concentration is high in the endoplasmic reticulum, and this compartment is tightly associated to the inclusion membrane (Derre, 2015). The unfolded protein response pathway is activated in infected cells (George, Omosun et al., 2016), a condition that might be sufficient to activate TG2 (Lee, Jeong et al., 2014). Indeed accumulation of cytoplasmic Ca^2+^ around the inclusion has been reported (Majeed, Krause et al., 1999), which might be enough to locally activate TG2.

Treatment of cells with a potent inhibitor of TG2, CP4d, reduced progeny ten-fold in primary epithelial cells (Fig. 2). We have shown that the inhibition of TG2 activity had two distinct effects on *C. trachomatis* developmental cycle. First, it reduced the ability for the bacteria to enter the cells (Fig. S2). *C. trachomatis* use multiple receptors and appear to hijack several entry pathways into epithelial cells (Ford, Nans et al., 2018). The positive role played by TG2 on bacterial entry, which could be exerted from its intracellular or extracellular location, remains to be studied in future work. One attractive candidate mechanism is PDGFR signaling, since it is implicated in *C. trachomatis* entry (Elwell, Ceesay et al., 2008) and it is sensitive to TG2 activity (Nurminskaya, Beazley et al., 2014). Second, the inclusion developed slower in CP4d-treated culture, indicating that TG2 is necessary for optimal bacterial growth. We discuss below the two mechanisms we uncovered that account for the link between TG2 activation and bacterial development, and place TG2 as a key regulator of bacterial access to glucose and its derivative UDP-GlcNAc.

We have shown that TG2 is required for the increase in transcription of *GLUT-1* and *GLUT-3* in infection in HeLa cells and in primary epithelial cells (Fig. 2). A similar observation was made in a mammary epithelial cell line (Kumar et al., 2014). In that case, it was shown that TG2 constitutively activated NF-κB expression, promoting the expression of the hypoxia-inducible factor (HIF)-1α, which in turn promoted *GLUT-1* transcription. A similar mechanism may take place in the case of *C. trachomatis* infection, since HIF-1 is also increased (Sharma, Machuy et al., 2011). Glucose is an essential metabolite for *C. trachomatis* development and preventing the transcription of *GLUT-1* and *GLUT-3* by siRNA led to a two-fold decrease in progeny in HeLa cells (Wang et al., 2017). This result shows that glucose import can become limiting for bacterial growth, and thus that the control exerted by TG2 on the expression of glucose transporters accounts, at least in part, for the need for this enzyme for optimal bacterial growth. This conclusion is supported by our observation that glucose becomes limiting faster for bacterial progeny in MEFs lacking TG2 than in wild-type cells. Thus, our study places TG2 as a key regulator for glucose import in infected cells.

Our proteomic approach identified 62 candidate TG2 substrates in *C. trachomatis* infection. Six of those were already known substrates of TG2, validating our analysis. We further demonstrated that GFPT was a substrate of TG2 and identified Q328 as a prominent transamidation site. The identification of which amine donor(s) are cross-linked to GFPT is left to future investigations, as several small amines might serve as substrates, making identification by mass spectrometry very challenging. We showed that GFPT activity was enhanced upon transamidation by TG2. This resulted in an increase in *O*-GlcNAcylation, since this post-translational modification of proteins is directly dependent on the concentration of UDP-GlcNAc (Jóźwiak, Forma et al., 2014, Love & Hanover, 2005). GFPT acts as a tetramer and is negatively regulated by several post-translational modifications (Chang, Su et al., 2000, Zibrova, Vandermoere et al., 2017) and by UDP-GlcNAc (Assrir, Richez et al., 2014). Transamidation on Gln328 could interfere with these down-regulation mechanisms and thereby unleash the HBP.

Interestingly the positive correlation between TG2 activation and *O*-GlcNAcylation does not hold true in infected cells and our data indicate that this is due to hexosamines being consumed by the infection. Our observation that silencing GFPT expression increases bacterial diameter strongly supports the hypothesis that UDP-GlcNAc, or an intermediate along the HBP, is hijacked into the inclusion to fuel bacterial division, possibly by feeding peptidoglycan synthesis. One recent publication showed that UDP-GlcNAc is also used during infection to post-translationally modify the intermediate filament vimentin and this also could contribute to significant UDP-GlcNAc consumption in *C. trachomatis* infection (Tarbet, Dolat et al., 2018).

The role played by TG2 in viral or microbial infections is raising increasing interest. Like in the case of *C. trachomatis* infection, genetic or pharmacological inhibition of TG2 led to a marked reduction in *Mycobacterium tuberculosis* replicative capacity. However, the mechanism involved might be different, since the data suggest that reduced replication in macrophages lacking TG2 is due to the impairment of autophagy homeostasis (Palucci, Matic et al., 2017). *M. tuberculosis* relies largely on lipids and fatty acids as energy source, and glucose availability might not be limiting in this case (Russell, VanderVen et al., 2010). Still, up-regulation of glucose transporter was also described in a mouse model of *M. tuberculosis* infection (Shi, Salamon et al., 2015), thus the involvement of TG2 in metabolism regulation in this context remains to be investigated.

There are multiple examples of host manipulation by pathogens that shed light on fundamental cellular processes. Here our work revealed an unsuspected regulation of the HBP by TG2. This discovery has important implications. Like other post-translational modifications, protein *O*-GlcNAcylation dramatically alters the fate and function of target proteins. In particular transcription factors are modified by *O*-GlcNAcylation, which implicates this modification in transcriptional regulation (Jackson & Tjian, 1988). Physiologically, disruption of *O*-GlcNAcylation homeostasis has been implicated in the pathogenesis of many human diseases, which include cancer, diabetes and neurodegeneration (Yang & Qian, 2017). The link between TG2 and *O*-GlcNAcylation means that TG2 activation is expected to have broad transcriptional consequences. In particular TG2 is activated in many cancers and future investigation is required to determine the contribution of the TG2/GFPT activation axis to tumorigenesis.

## MATERIAL AND METHODS

### Cells and bacteria

HeLa cells (ATCC) and mouse embryonic fibroblasts (MEFs) isolated from KO (TG2^−/−^) or WT (TG2^+/+^) C57B6 mice, kindly supplied by Dr. M. Piacentini (University of Rome), were grown in Dulbecco’s modified Eagle’s medium with Glutamax (DMEM, Invitrogen), supplemented with 10 % (v/v) heat-inactivated fetal bovine serum (FBS) (D’Eletto, Farrace et al., 2009). Primary cells used for experiments displayed in Fig. 2 were isolated from human fallopian tubes and maintained in culture as previously described (Roth et al., 2010). Other primary cells were isolated from endocervix biopsies of female patients and were cultivated in keratinocyte-SFM medium (Thermo Fisher Scientific) containing 50 mg.L^−1^ of bovine pituitary extract (Thermo Fisher Scientific) and 5 μg.L^−1^ of epidermal growth factor (EGF) human recombinant (Thermo Fisher Scientific) (Wu *et al*, in preparation). ES-2 cells were grown in DMEM supplemented with 10 % FBS, 2 mM L-glutamine (Sigma #59202C), 1x MEM non-essential amino acids (Sigma M7145) and 1 mM sodium pyruvate. PEO4 cells were grown in RPMI supplemented with 10 % FBS, 2 mM L-glutamine (Sigma #59202C), 1x MEM non-essential amino acids (Sigma M7145), 1 mM sodium pyruvate and 1x insulin (Sigma I9278). All cell cultures were maintained at 37 °C, in 5 % CO2 atmosphere and were routinely tested for mycoplasma using the standard PCR method. *C. trachomatis* serovar LGV L2 strain 434 and serovar D/UW-3/CX (ATCC), GFP-expressing L2 (L2^IncD^GFP) or *C. muridarum* MoPn (for *in* vivo experiments) were propagated on HeLa cells, purified on density gradients as previously described and stored at −80 °C (Scidmore, 2005, Vromman, Laverriere et al., 2014).

### siRNA treatment

For siRNA experiments, 50 000 cells were plated in a 24-well plate and immediately mixed with Lipofectamine RNAiMAX (Invitrogen) following the manufacturer’s recommendation, using 10 nM of siRNA (Table S2). For RB size assessment, 300 000 cells were plated in a 6-well plate. For electron microscopy experiments, 1.5 million cells were plated in a 25-cm^2^ flask. For GFPT1 activity assay, 1 million cells were plated in a 10-cm diameter dish. The culture medium was changed the next day and experiments (infection or treatment with ionomycin) were performed two days post treatment with the siRNA.

### GFPT1 purification

Recombinant human GFPT1 (rhGFPT1) with an internal 6-His tag was produced from a plasmid pET28-rhGFPT1-6His in *E. coli* Rosetta (DE3) GlmS::Tc kindly given by Dr. Badet-Denisot (Centre de Recherche de Gif, France) (Li, Roux et al., 2007). The mutated form rhGFPT1 Q58N, rhGFPT1 Q328N and rhGFPT1 Q546N were obtained using QuikChange technology (Agilent) on the plasmid pET28-rhGFPT1-6His, with primers listed in Table S2, following the manufacturer instructions, and transformed in *E. coli* Rosetta (DE3) GlmS::Tc. One litre of culture in 2YT medium supplemented with tetracycline (8 μg.mL^−1^, Sigma), kanamycin (50 μg.mL^−1^, Sigma), chloramphenicol (15 μg.mL^−1^, Sigma) and glucosamine (GlcNH_2_, 2 mg.mL^−1^, Sigma) was incubated with agitation at 37 °C until OD_600_ reached 0.5. Protein expression was induced by addition of 0.5 mM of isopropyl β-d-thiogalactopyranoside (Sigma) at 25 °C for 24 h before being harvested. The cell pellets were resuspended in lysis buffer (16.27 mM Na2HPO4 and 3.73 mM NaH2PO4 pH 7.5, 200 mM NaCl, 20 mM imidazole, 2 mM tris(2-carboxyethyl)phosphine hydrochloride (TCEP), 10 % glycerol, 1 mM fructose-6-P, Roche EDTA-free protease inhibitor cocktail) and disrupted by sonication. The recombinant protein was purified by incubation with Qiagen Ni-NTA agarose beads (Qiagen) for 1 h followed by three washing steps with the lysis buffer before elution with lysis buffer containing increasing concentrations of imidazole: 30 mM, 100 mM, 175 mM and finally 500 mM of imizadole. The fractions containing the protein were dialyzed against lysis buffer with 20 mM HEPES replacing the phosphate buffer before storage at −80 °C

### TG2 activity assay

#### In vivo

Cells plated the day before (100 000 cells/well) were infected with *C. trachomatis* serovar LGV L2 at a MOI of 1, and 0.5 μM biotin pentylamine (BP) (Thermo Fisher Scientific) was added after 24 h. In some experiments, cells were pre-incubated for 2 h with 40 μM CP4d or DMSO as control before addition of bacteria. CP4d inhibits the transamidase activity of TG2 (K_i_ = 174 nM) and favours its closed conformation (Caron et al., 2012). For experiments with ionomycin (Sigma), cells pre-treated with siRNA for 48 h or infected as described above for 24 h were treated with Ionomycin and 0.5 μM BP for 6 h.

At the end of the indicated incubation time, cells were lysed using 8 M urea buffer (30 mM Tris, 150 mM NaCl, 8 M urea, 1 % SDS, pH=8.0) and samples subjected to Western Blot.

#### In vitro

1 mU of transglutaminase from guinea pig liver (Sigma) was incubated at 37 °C for 15 min or 3 h with 5 μg of rhGFPT1, rhGFPT1 Q58N or rhGFPT1 Q328N and 1 mM of BP in 20 mM HEPES buffer pH 7.5, 200 mM NaCl, 2 mM TCEP, 10 % glycerol, 1 mM fructose-6-phosphate, 10 mM CaCl2. The reaction was stopped by adding ethylene-bis(oxyethylenenitrilo)tetraacetic acid (EGTA) at a final concentration of 20 mM and boiling the samples 5 min at 95 °C.

### Streptavidin-precipitation of TG2 targets

Twenty million HeLa cells were seeded in a 163-cm^2^ flask. One day later, the cells were infected or not with *C. trachomatis* serovar LGV L2 at a MOI = 1. Two hours post infection (hpi) the culture medium was changed and 40 μM CP4d or DMSO was added. BP was added to the culture medium at 0.5 mM 24 h post treatment (infection or not) and cells were lysed 46 hpi directly in the well using 8 M urea buffer. DNA was disrupted by sonication and a dialysis was performed against 2 M urea buffer (Tris 30 mM, NaCl 150 mM, 2 M urea, 1 % SDS, pH=8.0). Samples were incubated overnight at 4 °C with streptavidin-agarose beads (Sigma). After 3 washes with 2 M urea buffer and 3 washes with phosphate-buffered saline (PBS), proteins precipitated on the beads were eluted using Laemmli’s buffer containing dithiothreitol (Sigma) boiled 5 min at 95 °C. Samples were then analyzed by Western blot or by mass spectrometry.

### SDS-PAGE and Western blot

Proteins were subjected to sodium dodecyl-sulfate polyacrylamide gel electrophoresis (SDS-PAGE) and transferred to a polyvinylidene difluoride (PVDF) membrane, which was blocked with 1 × PBS containing 5 % bovine serum albumin (BSA, for biotin revelation only) or milk and 0.01 % Tween 20. The membranes were then immunoblotted with primary antibodies diluted in 1 × PBS containing 5 % milk and 0.01 % Tween 20. For analyzing the TG2 activity assay, biotin incorporation was revealed using streptavidin conjugated to HRP (Sigma). Primary antibodies used in the western blots were the mouse clone 7D2 against TG2 (#ABIN1109303, Covalab), rabbit anti-serum against GFPT (kindly given by Dr. C. Weigert, University of Tübingen, Germany), mouse clone RL2 against *O*-GlcNAcylation (#MA1-072 Thermo Fisher Scientific) and mouse clone AC-74 against β-actin (#A5441 Sigma). Secondary antibodies were anti-mouse-HRP (GE Healthcare) or anti-rabbit-HRP (Invitrogen) conjugated antibodies. Blots were developed using the Western Lightning Chemiluminescence Reagent (GE Healthcare).

### Ovarian cancer cohort and statistical analysis

Expression data for *TGM2, GLUT-1* and *GLUT-3* in 265 high grade serous ovarian cancers from Edinburgh were available from previous transcriptomic studies of ovarian cancer (Hollis, Churchman et al., 2019). Per-sample expression was calculated as the mean expression of probe-sets informative for each gene. Expression comparisons were performed using Spearman’s rank correlation test.

### Mass Spectrometry

#### In solution protein digestion

Samples were prepared in triplicate. For streptavidin-precipitation of TG2 targets samples, tryptic digestion was performed by enhanced filter-aided sample preparation. All steps were done in-filter. Briefly, samples were reduced (50 mM TCEP, 30 minutes at room temperature) and alkylated (50 mM iodoacetamide, 1 h at room temperature in the dark). Then, proteins were incubated overnight at 37 °C with 500bng trypsin (Trypsin Gold Mass Spectrometry Grade, Promega). Peptides were recovered by centrifugation.

After TG2/GFPT1 reactions *in vitro* samples were diluted in a large excess of 8 M urea / 100 mM Tris HCl pH 8.5 buffer and then, as previously described, reduced (5 mM TCEP, 30 minutes at room temperature) and alkylated (10 mM iodoacetamide, 30 minutes at room temperature in the dark). Proteins were first digested for 5 h at 37 °C with 500 ng rLys-C Mass Spec Grade (Promega, Madison, WI, USA) before being diluted 4-fold with 100 mM Tris HCl pH 8.5 to reach a concentration below 2 M urea. Samples were then incubated overnight at 37 °C with 500 ng Sequencing Grade Modified Trypsin (Promega, Madison, WI, USA). To achieve the complete digestion of the peptides, a second incubation with the same amount of trypsin (5 h at 37 °C) was performed. Digestion was stopped by adding formic acid to 5 % final concentration and peptides were desalted and concentrated on Sep-Pak C_18_SPE cartridge (Waters, Milford, MA, USA) according to manufacturer instructions.

#### Mass spectrometry analysis

Tryptic peptides were analyzed on a Q Exactive Plus instrument (Thermo Fisher Scientific, Bremen) coupled with an EASY nLC 1 000 or 1 200 chromatography system (Thermo Fisher Scientific, Bremen). Sample was loaded on an in-house packed 50 cm nano-HPLC column (75 μm inner diameter) with C_18_ resin (1.9 μm particles, 100 Å pore size, Reprosil-Pur Basic C_18_-HD resin, Dr. Maisch GmbH, Ammerbuch-Entringen, Germany) and equilibrated in 98 % solvent A (H_2_O, 0.1 % FA) and 2 % solvent B (ACN, 0.1 % FA). 120 or 180 min gradient of solvent B at 250 nL.min^−1^ flow rates were applied to separated peptides. The instrument method for the Q Exactive Plus was set up in DDA mode (Data Dependent Acquisition). After a survey scan in the Orbitrap (resolution 70 000), the 10 most intense precursor ions were selected for HCD fragmentation with a normalized collision energy set up to 28. Charge state screening was enabled, and precursors with unknown charge state or a charge state of 1 and > 7 were excluded. Dynamic exclusion was enabled for 35 or 45 seconds respectively.

#### Data processing

Data were searched using Andromeda with MaxQuant software 1.4.1.2 or 1.5.3.8 version against respectively a *Chlamydia trachomatis* Uniprot reference proteome database concatenated with Homo sapiens Uniprot reference proteome database. Data were also searched against usual known mass spectrometry contaminants and reversed sequences of all entries or an *E. coli* K12 Uniprot reference proteome database concatenated with rhGFPT1 and gpTGase proteins (Tyanova, Temu et al., 2016). Andromeda searches were performed choosing trypsin as specific enzyme with a maximum number of two missed cleavages. Possible modifications included carbamidomethylation (Cys, fixed), oxidation (Met, variable), N-ter acetylation (variable) and BP (Gln, variable). The mass tolerance in MS was set to 20 ppm for the first search then 6 ppm for the main search and 10 ppm for the MS/MS. Maximum peptide charge was set to seven and five amino acids were required as minimum peptide length. The “match between runs” feature was applied between replicates with a maximal retention time window of 2 or 0.7 min. One unique peptide to the protein group was required for the protein identification. A false discovery rate (FDR) cutoff of 1 % was applied at the peptide and protein levels.

#### Data analysis

To validate the identification of the BP on the glutamine of modified peptides, spectra were manually inspected (or fragment assignments).

For the global quantification, output files from MaxQuant were used for protein quantification. Quantification was performed using the XIC-based LFQ algorithm with the Fast LFQ mode as described previously (Cox, Hein et al., 2014). Unique and razor peptides, included modified peptides, with at least 2 ratio count were accepted for quantification.

For pairwise comparisons, proteins identified in the reverse and contaminant databases and proteins only identified by site were first discarded from the list. Then, proteins exhibiting fewer than 2 LFQ values in at least one condition were discarded from the list to avoid misidentified proteins. After log2 transformation of the leftover proteins, LFQ values were normalized by median centering within conditions (normalizeD function of the R package DAPAR (Wieczorek, Combes et al., 2017)). Remaining proteins without any LFQ value in one of both conditions have been considered as proteins quantitatively present in a condition and absent in another. They have therefore been set aside and considered as differentially abundant proteins. Next, missing values were imputed using the imp.norm function of the R package norm. Proteins with a foldchange under 2 have been considered not significantly differentially abundant. Statistical testing of the remaining proteins (having a foldchange over 2) was conducted using a limma t-test thanks to the R package limma (Ritchie, Phipson et al., 2015). An adaptive Benjamini-Hochberg procedure was applied on the resulting p-values thanks to the function adjust.p of R package cp4p using the robust method of Pounds and Cheng to estimate the proportion of true null hypotheses among the set of statistical tests (Pounds & Cheng, 2006). The proteins associated to an adjusted p-value inferior to an FDR level of 1% have been considered as significantly differentially abundant proteins.

### Adhesion assay

Adhesion assays were performed as described previously (Vromman et al., 2014). In brief, MEFs cells plated in 24-well plate the day before (100 000 cells/well) were pre-cooled 30 min at 4 °C and then were incubated for 4 h at 4 °C with L2^IncD^GFP bacteria at a MOI = 10, sonicated prior to infection in order to disrupt bacterial aggregates. Then cells were washed gently with PBS and detached using 0.5 mM EDTA in PBS. Samples were fixed 30 min in 2 % PFA, washed with PBS and analyzed using flow cytometry.

### Bacterial entry assessment

Entry experiments were performed as described previously (Vromman et al., 2014). In brief, MEFs cells plated on coverslips in 24-well plate the day before (100 000 cells/well) were pre-cooled 30 min at 4 °C and then incubated for 45 min at 4 °C with L2^IncD^GFP bacteria at a MOI = 10, sonicated prior to infection in order to disrupt bacterial aggregates. Then pre-warmed medium was added and coverslips were incubated at 37 °C before being fixed at different time points in 4 % PFA for 20 min. Extracellular bacteria were stained with a mouse anti-MOMP-LPS (Argene # 11-114) antibody followed with Cy5-conjugated anti-mouse (Amersham Biosciences) secondary antibody. The dilutions were made in PBS containing 3 % of BSA. DNA was stained using 0.5 μg.mL^−1^ of Hoechst 33342 (Thermo Fisher Scientific) added in the secondary antibody solution. Images were acquired on an Axio observer Z1 microscope equipped with an ApoTomemodule (Zeiss, Germany) and a 63× Apochromat lens. Images were taken with an ORCAflash4.OLT camera (Hamamatsu, Japan) using the software Zen.

### Progeny assay

For glucose privation tests on MEFs cells, 140 000 cells per well were seeded in a glucose-free DMEM (Invitrogen) supplemented with 10 % FBS. The following day, the medium was replaced with glucose-free DMEM supplemented with 10 % FBS and the indicated concentration of glucose (Sigma). The next day cells were infected with L2^IncD^GFP bacteria at a MOI = 0.2. For progeny assays on HeLa cells, primary cells or MEFs cells, 100 000 cells were seeded in a 24-well plate. The next day cells were pre-treated with CP4d (or DMSO) for 2 h and infected with L2^IncD^GFP bacteria at a MOI = 0.15. Cells treated with siRNA for 48 h were directly infected with L2^IncD^GFP bacteria at a MOI = 0.2. Thirty hpi, cells were detached and fixed in 2 % PFA in PBS prior to flow cytometry analysis in order to evaluate the first round of infection. In duplicate wells, cells were detached, lysed using glass beads and the supernatant was used to infect new untreated cells (or WT cells in the case of MEFs) plated the day before (100 000 cells/well in a 24-well plate), in serial dilution. The next day, 3 wells per condition with an infection lower than 30 % (checked by microscopy) were detached and fixed as described above, before analysis by flow cytometry and determination of the bacterial titer. Acquisition was performed using a CytoFLEX S (Beckman Coulter) and 50 000 events per sample were acquired and then analyzed using FlowJo (version 10.0.7).

### Infection in mice

Female TG2^−/−^ and TG2^+/+^ KO mice were kindly provided by Dr. C. Papista (INSERM UMR970, Centre de Recherche Cardiovasculaire, Paris) and maintained in the animal facility of the Institut Pasteur, Paris. All animals are treated with 2.5 mg of medroxyprogesterone (Depo-provera-SC®, Pfizer) 7 days prior to infection to synchronize the menstrual cycle. Mice were intravaginally inoculated with *C. muridarum*, 10^5^ IFU per animal. Twenty-five days after infection, animals were sacrificed and the organs excised including cervix, uterine horn and oviduct. The bacterial burden in the excised organs, in the right part of the upper genital tract was measured by qPCR after DNA extraction using the DNeasy Blood and Tissue Kit (Qiagen). The left part of the upper genital tract was excised and rinsed into PBS for the morphological observation. Hydrosalpinx score was determined as described (Peng, Lu et al., 2011).

### RT-qPCR and qPCR

One hundred twenty-five thousand cells in a 24-well plate were infected or not with *C. trachomatis* serovar LGV L2 at a MOI = 1. Cells were treated with CP4d (40 μM) or DMSO 2 hpi, or doxycycline (62.5 ng.mL^−1^, Sigma) 24 hpi. Total RNAs were isolated 24 or 48 hpi with the RNeasy Mini Kit (Qiagen) with DNase treatment (DNase I, Roche). RNA concentrations were determined with a spectrophotometer NanoDrop (Thermo Fisher Scientific) and normalized to equal contents. Reverse transcription (RT) was performed using the M-MLV Reverse Transcriptase (Promega) and quantitative PCR (qPCR) undertaken on the complementary DNA (cDNA) with LightCycler 480 system using LightCycler 480 SYBR Green Master I (Roche). For the experiments displayed in Fig. S4, a duplicate well was used to extract genomic DNA (gDNA) of each time point using the DNeasy Blood and Tissue Kit (Qiagen). Data were analyzed using the ΔΔCt method with the *actin* gene as a control gene (Schmittgen & Livak, 2008).

### GFPT activity assay

HeLa cells treated with ionomycin (4 μM) or DMSO for 6 h were detached in lysis buffer containing 0.05 M Tris, 0.15 M NaCl, 5 % glycerol, 0.5% NP-4, protease inhibitor cocktail EDTA-free (Roche), pH 7.5. After lysis at 4 °C, NP-40 concentration was reduced by addition of an excess of reaction buffer (0.05 M Tris, 0.15 M NaCl, 5 % glycerol, protease inhibitor cocktail EDTA-free, pH 7.5) and cell debris were eliminated by centrifugation. Cell lysates were incubated 45 min at 37 °C with 0.6 mg.mL^−1^ fructose-6-phosphate (Sigma) and 0.6 mg.mL^−1^ glutamine (Sigma). The reaction was stopped by incubating the samples for 5 min at 100 °C and precipitates were removed by centrifugation. Analysis of the samples was performed by high performance anion exchange chromatography (HPAEC, Dionex, model ISC3000) on a CarboPAC-PA1 column (3.2 × 250 mm, Dionex) using 100 mM NaOH, and 720 mM NaOAc in 100 mM NaOH, as eluent A and B, respectively. The column was pre-equilibrated for 20 min in 98 % A + 2 % B. Following sample injection, a gradient run (flow rate 1 mL.min^−1^) was performed as follows: 0–2 min, isocratic step (98 % A + 2 % B), 2–15 min 98 % A + 2 % B – 80 % A + 20 % B, 15–20 min 80 % A + 20 % B – 57 % A + 43 % B, 20–22 min 57 % A + 43 % B – 100 % B, and 22–25 min 100 % B. Samples were detected on a pulsed electrochemical detector.

### Bacterial size measurement

HeLa cells in a 6-well plate, treated with siRNA as described above, were infected every two hours with L2^IncD^GFP bacteria at a MOI = 0.3. The next day, all wells were detached and fixed simultaneously in 2 % PFA and 2.5 % glutaraldehyde (Sigma) in PBS. After 25 min, cells were broken using glass beads, vortexed and syringed (3 times, using 1 mL 26GA × 3/8-inch syringes). Samples were then analyzed by flow cytometry. An exponential culture of *E. coli*, purified *C. trachomatis* elementary bodies (i.e. the infectious non-replicative form of the bacterium) and non-infected cells prepared the same way were used to gate successively for particles of equal size or smaller than *E. coli* (thereby excluding non-broken cells), larger than elementary bodies (thereby excluding non-dividing bacteria), and with positive green fluorescence (thereby excluding cell debris). The forward-scattered light (FSC-A) was used to compare bacterial diameters. Each data point represents the mean of at least 400 gated events.

### Electron microscopy

One million five hundred thousand cells were transfected with siRNA and infected two days later. Cells were fixed 30 hpi with 2.5 % glutaraldehyde (v/v) (Electron Microscopy Sciences) in 0.1 M cacodylate buffer pH 7.4, for 1 h at room temperature. After several washes in cacodylate they were post-fixed with 1 % osmium tetroxide (w/v) in cacodylate for 1 h. After several washes with water the cells were progressively dehydrated with increasing concentrations of ethanol from 25 % to 100 %. The cells were then gradually embedded in epoxy resin. After overnight polymerization at 60 °C, 50 to 70 nm thin sections were cut in an ultra-microtome (Ultracut, Leica) and cells were imaged after post-staining with uranyl acetate and lead citrate in a T12-FEI transmission EM operated at 120kV.

## Supporting information

All Suppl material

## ACKNOWLEDGEMENTS

We thank Anke Hellberg for technical assistance, Dr Christina Papista for providing mice, Mauro Piacentini for MEFs, Dr Denise Badet-Denisot for the rhGFPT1 plasmid and for advice, Dr Cora Weigert for anti-GFPT antibodies, Vishu Aimanianda Bopaiah for help with HPAEC. This work was supported by an ERC Starting Grant (NUChLEAR N°282046), the Institut Pasteur (GFP-LIMNEC METINF) and the Centre National de la Recherche Scientifique. BM was funded by the Ministère de l’Education Nationale, de la Recherche et de la Technologie and by Cancéropole Ile-de-France.

---

Figure S1. **Live bacteria induce TG2 activation in HeLa and MEFs cells**. – Related to Figure 1. Before addition to HeLa cells the bacterial inoculum was either left untreated (MOI = 1), or filtered through of a 0.22 μM filter, or incubated for 30 min at 65 °C to kill the bacteria. The inoculum was applied to HeLa cells in the presence or not of BP and 48 h later whole cell lysates were prepared. After separation by SDS-PAGE, proteins were transferred to a membrane and BP incorporation was revealed with HRP-conjugated streptavidin.

Figure S2. **TG2 is beneficial for bacterial development**. – Related to Figure 2. A – TG2^+/+^ and TG2^−/−^ MEFs were infected with L2^incD^GFP at MOI=0.15. Thirty h later the cells were disrupted and bacterial titers were determined by re-infecting fresh TG2 ^+/+^ cells as described in the methods. The mean ± SD of five independent experiments are shown. B – To measure bacterial adhesion TG2^+/+^ and TG2^−/−^ MEFs were incubated at 4 °C for 4 h with L2^incD^GFP at MOI=10 before being washed and fixed as described in the methods. The mean ± SD of three independent experiments are shown. C – TG2^+/+^ and TG2^−/−^ MEFs were infected with L2^incD^GFP at MOI=10 and fixed at the indicated time. Extracellular bacteria were differentially labeled as described in the methods. The mean ± SD of three independent experiments are shown. Asterisks indicate two significantly different values (*: P < 0.05; **: P < 0.01, Student’s paired t-test).

Figure S3. **Infection by *C. muridarum* activates TG2**. – Related to Figure 2. Whole cell lysates were prepared with HeLa cells infected or not for 48 h with *C. muridarum* (MOI=1) in the presence or not of BP. Cell lysates were run on SDS-PAGE, proteins were transferred to a membrane and BP incorporation was revealed with HRP-conjugated streptavidin.

Figure S4. **Reduction of bacterial load does not account for the loss of induction of *GLUT-1* and *GLUT-3* transcription upon CP4d treatment**. Related to Figure 3. HeLa cells were infected or not with *C. trachomatis* (MOI=1) in duplicate per condition. Two hpi CP4d (40 μM) or 24 hpi doxycycline (62.5 ng.mL^−1^) were added to the culture medium. Forty-eight hpi DNA were extracted from one well to measure bacterial load, and RNA were extracted from the duplicate well. Bacterial gDNA (16S RNA) measured by real-time RT-PCR was normalized to host gDNA (*actin* gene) and is expressed relative to the infected non-treated culture. *GLUT-1* and *GLUT-3* transcripts were measured by real-time RT-PCR and normalized to *actin* with the ΔΔCt method as in Fig. 3A. The experiment was performed in duplicate and repeated four times.

Figure S5. **Measure of glucosamine-6-P production by GFPT by HPAEC-PAD**. Related to Figure 4. The three top panels show the retention times of the different sugars used or produced during the reaction when analyzed by HPAEC-PAD. Note that glutamine and glutamate are not retained by this column. A cell lysate without addition of fructose-6-P or glutamine is also shown. The bottom two panels display an enlargement of the output of the reaction when control or ionomycin treated cell lysates were used. The arrows point to the glucosamine-6-P peak. Fructose-6-P was not entirely consumed by the reaction. Formation of glucose-6-P was also observed, possibly by GFPT isomerase activity or by another cellular enzyme present in the lysate such as glucose-6-P isomerase.

Figure S6. **Gating steps for the examination of the size of replicative *C. trachomatis* by flow cytometry**. Related to Figure 6. Exponential cultures of *E. coli*, density-gradient purified elementary bodies, non-infected cells, and cells infected with L2^incD^GFP at MOI=0.3 for different time points (indicated) were fixed and treated as described in the methods section. The top dot plots describe how the first three samples were used to gate the region of interest that contains mostly replicative *C. trachomatis* (excluded regions are hatched). An increase in the number of replicative bacteria, and a decrease in their size (FSC-A value) were observed when increasing infection time.

Figure S7. **GFPT silencing affects bacterial division**. Related to Figure 6. HeLa cells treated for 48 h with siRNA targeting GFPT1 or not (siCTRL) were infected with *C. trachomatis* (MOI = 1), fixed 30 hpi and processed for transmission electron microscopy. Lines show example of measured RB diameters. Scale bar = 600 nm. RB diameters were measured using ImageJ. The mean value is indicated. Asterisks indicate two significantly different values (**: P < 0.01, Student’s paired t-test).

Table S1. **Candidate TG2 substrates in *C. trachomatis* infected cells** The list displays streptavidin-bound proteins identified by mass spectrometry in the infected samples only in the presence of BP (first 37 entries) or more abundant in the presence of BP than in its absence (last 25 entries, Log2([Mean intensity with BP]/[Mean intensity without BP]) and p-values are shown).

Table S2. **Primers used for siRNA, qPCR and mutagenesis**

