## Supplementary material for "Infection-driven activation of transglutaminase 2 boosts glucose uptake and hexosamine biosynthesis": All Suppl material

Figure S1

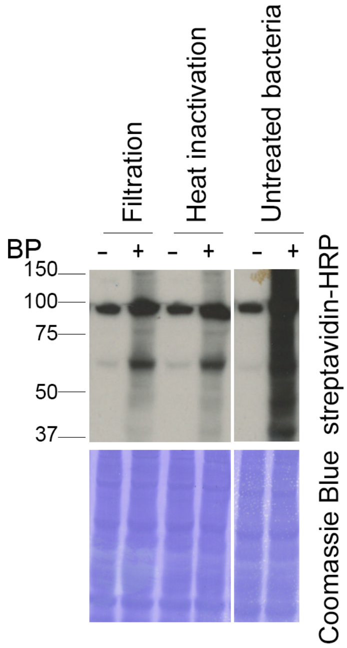

Filtration
Heat inactivation
Untreated bacteria
BP
- + - + - +
150
100
75
50
37
streptavidin-HRP
Coomassie Blue

Figure S2

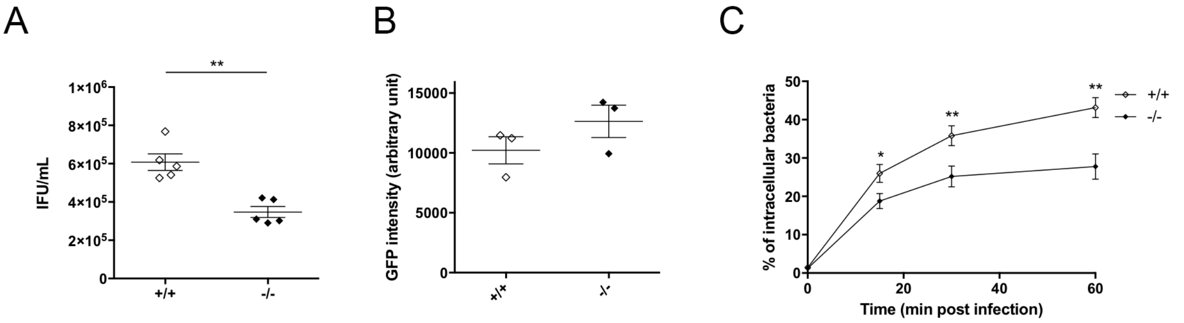

A
**
IFU/mL
1×10^6
8×10^5
6×10^5
4×10^5
2×10^5
0
+/+
-/-
B
GFP intensity (arbitrary unit)
15000
10000
5000
0
+/+
-/-
C
% of intracellular bacteria
50
40
30
20
10
0
0
20
40
60
Time (min post infection)
*
**
**
+/+
-/-

Figure S3

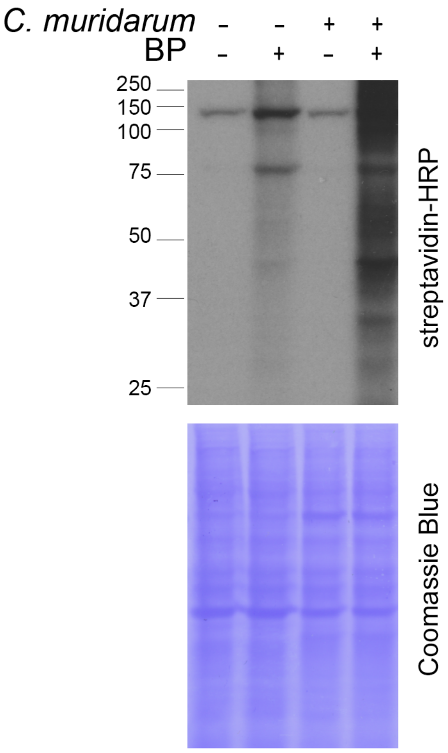
C. muridarum
BP
250
150
100
75
50
37
25
streptavidin-HRP
Coomassie Blue

### Figure S4

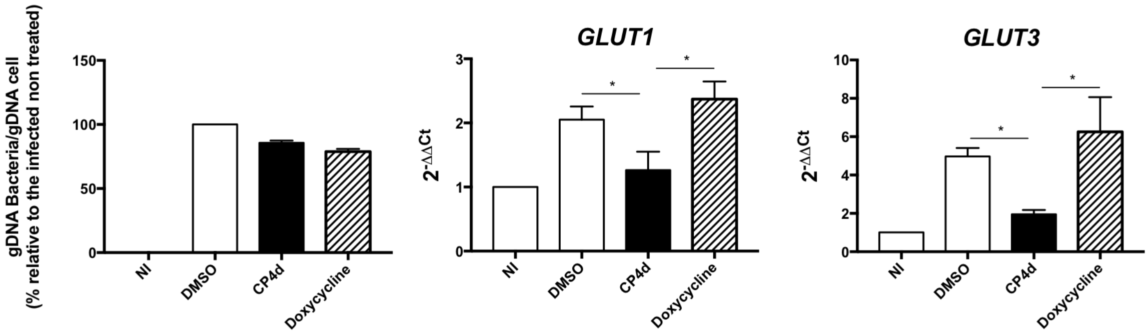
gDNA Bacteria/gDNA cell
(% relative to the infected non treated)
150
100
50
0
NI
DMSO
CP4d
Doxycycline
GLUT1
$2^{-\Delta\Delta Ct}$
3
2
1
0
*
*
NI
DMSO
CP4d
Doxycycline
GLUT3
$2^{-\Delta\Delta Ct}$
10
8
6
4
2
0
*
*
NI
DMSO
CP4d
Doxycycline

Figure S5

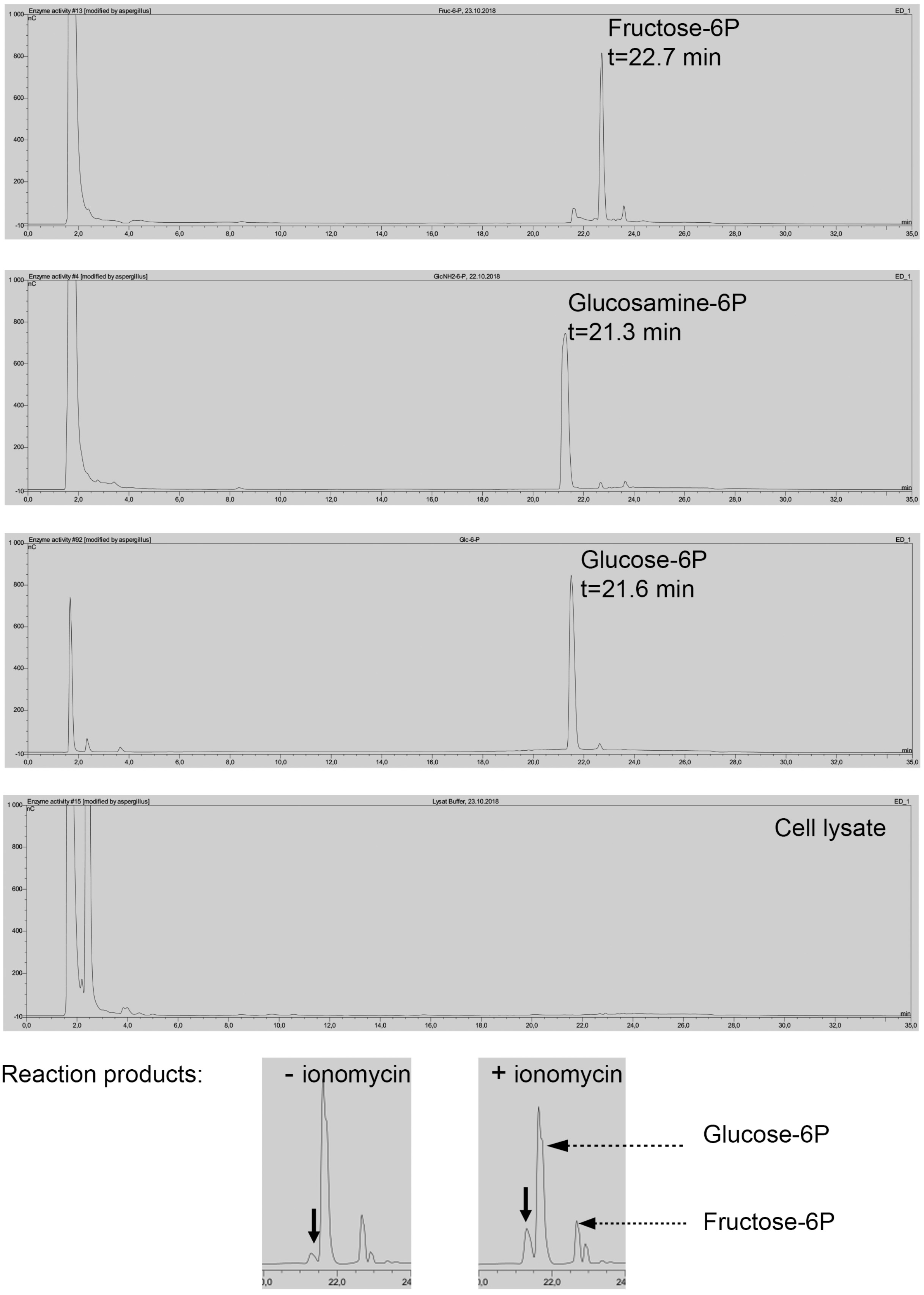

Fructose-6P
t=22.7 min
Glucosamine-6P
t=21.3 min
Glucose-6P
t=21.6 min
Cell lysate
Reaction products:
- ionomycin
+ ionomycin
Glucose-6P
Fructose-6P

Figure S6

1 – Gating on *E. coli* 2 – Exclusion of non-dividing bacteria (elementary bodies) 3 – Exclusion of cell debris

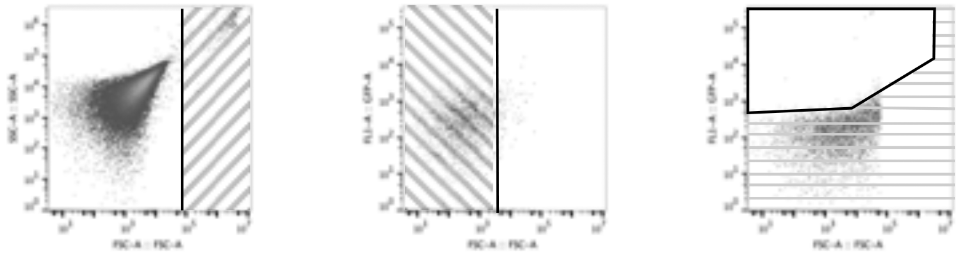

4 - Infected sample analysis

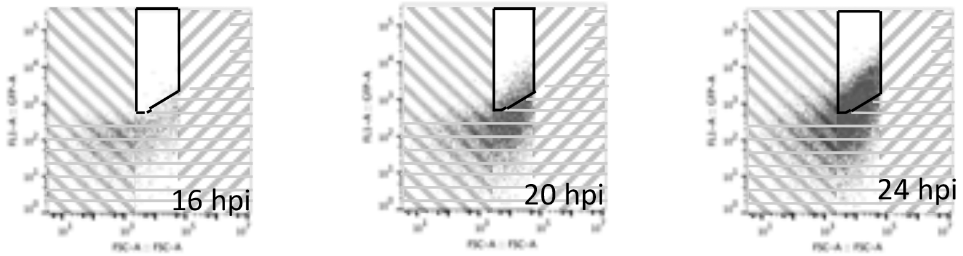


| Infection time (h) | 16 | 20 | 24 |
|---|---|---|---|
| Number of events gated | 53 | 1300 | 5596 |
| FSC-A Geometric Mean | 24688 | 18135 | 14445 |

Figure S7

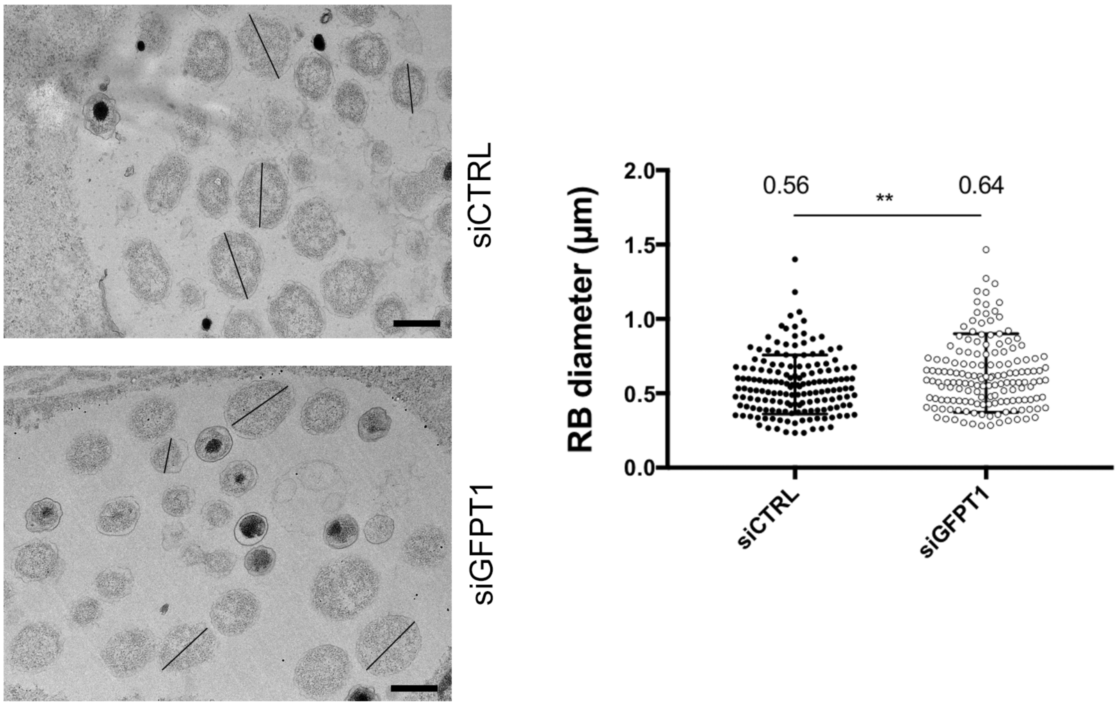
siCTRL
siGFPT1
RB diameter (µm)
2.0
1.5
1.0
0.5
0.0
0.56
0.64
**
siCTRL
siGFPT1

**Table S1: Candidate TG2 substrates in *C. trachomatis* infected cells**

The list displays streptavidin-bound proteins identified by mass spectrometry in the infected samples only in the presence of BP (first 37 entries) or more abundant in the presence of BP than in its absence (last 25 entries, Log2([Mean intensity with BP]/[Mean intensity without BP]) and p-values are shown).

| UniProt ID | Protein | Gene | Protein description | log2 | P-value |
|---|---|---|---|---|---|
| Q8NE71-2 | ABCF1 | ABCF1 | ATP-binding cassette sub-family F member 1 | NA | NA |
| O95831-3 | AIFM1 | AIFM1 | Apoptosis-inducing factor 1 | NA | NA |
| P50995-2 | ANX11 | ANXA11 | Annexin A11 | NA | NA |
| P84077 | ARF1 | ARF1 | ADP-ribosylation factor 1 | NA | NA |
| Q13510 | ASAH1 | ASAH1 | Acid ceramidase | NA | NA |
| Q12797-10 | ASPH | ASPH | Aspartyl/asparaginyl beta-hydroxylase | NA | NA |
| P36542-2 | ATPG | ATP5C1 | ATP synthase subunit gamma | NA | NA |
| O95816 | BAG2 | BAG2 | BAG family molecular chaperone regulator | NA | NA |
| Q13185 | CBX3 | CBX3 | Chromobox protein homolog 3 | NA | NA |
| Q9Y696 | CLIC4 | CLIC4 | Chloride intracellular channel protein 4 | NA | NA |
| Q16555-2 | DPYL2 | DPYSL2 | Dihydropyrimidinase-related protein | NA | NA |
| P55884 | EIF3B | EIF3B | Eukaryotic translation initiation factor 3 subunit B | NA | NA |
| P02751-4 | FINC | FN1 | Fibronectin | NA | NA |
| G3V1Q4 | G3V1Q4 | SEPT7 | Septin 7 | NA | NA |
| Q9HAV0 | GBB4 | GNB4 | Guanine nucleotide-binding protein subunit beta | NA | NA |
| O94808 | GFPT2 | GFPT2 | Glutamine--fructose-6-phosphate aminotransferase | NA | NA |
| P08754 | GNAI3 | GNAI3 | Guanine nucleotide-binding protein G(k) subunit alpha | NA | NA |
| H3BSW3 | H3BSW3 | APRT | Adenine phosphoribosyltransferase | NA | NA |
| P68871 | HBB | HBB | Hemoglobin subunit beta | NA | NA |
| A0A087WZW8 | IGKV3-11 | IGKV3-11 | Ig kappa chain C region | NA | NA |
| Q15181 | IPYR | PPA1 | Inorganic pyrophosphatase | NA | NA |
| J3QLE5 | J3QLE5 | SNRPN | Small nuclear ribonucleoprotein-associated protein N | NA | NA |
| K7ENG2 | K7ENG2 | U2AF2 | Splicing factor U2AF 65 kDa subunit | NA | NA |
| P17931 | LEG3 | LGALS3 | Galectin-3 | NA | NA |
| C9JIG9 | OSR1 | OSXR1 | Serine/threonine-protein kinase OSR1 | NA | NA |
| P17858 | PFKAL | PFKL | ATP-dependent 6-phosphofructokinase | NA | NA |
| E9PQ98 | PRMT1 | PRMT1 | Protein arginine N-methyltransferase 1 | NA | NA |
| P62195-2 | PRS8 | PSMC5 | 26S proteasome regulatory subunit 8 | NA | NA |
| O00487 | PSDE | PSMD14 | 26S proteasome non-ATPase regulatory subunit | NA | NA |
| Q16401-2 | PSMD5 | PSMD5 | 26S proteasome non-ATPase regulatory subunit 5 | NA | NA |
| Q06203 | PUR1 | PPAT | Amidophosphoribosyltransferase | NA | NA |
| P17812-2 | PYRG1 | CTPS1 | CTP synthase 1 | NA | NA |
| P43487-2 | RANG | RANBP1 | Ran-specific GTPase-activating protein | NA | NA |
| Q9P2E9-2 | RRBP1 | RRBP1 | Ribosome-binding protein 1 | NA | NA |
| P56192 | SYMC | MARS | Methionine--tRNA ligase | NA | NA |
| O95497 | VNN1 | VNN1 | Pantetheinase | NA | NA |

| | | | | | |
|---|---|---|---|---|---|
| E9PRD9 | VNN2 | VNN2 | Vascular non-inflammatory molecule 2 | NA | NA |
| P08243-2 | ASNS | ASNS | Asparagine synthetase | 7,62 | 1,81E-08 |
| P30508 | HLA-C | HLA-C | HLA class I histocompatibility antigen, Cw-12 alpha chain | 9,05 | 4,07E-08 |
| P35613-2 | BASI | BSG | Basigin | 2,33 | 1,85E-05 |
| P10909-4 | CLUS | CLU | Clusterin | 2,13 | 3,91E-05 |
| P49368 | TCPG | CCT3 | T-complex protein 1 subunit gamma | 1,83 | 3,99E-04 |
| O15427 | MOT4 | SLC16A3 | Monocarboxylate transporter 4 | 1,93 | 6,21E-04 |
| E9PLL6 | RPL27A | RPL271 | 60S ribosomal protein L27a | 1,36 | 9,12E-04 |
| Q15233 | NONO | NONO | Non-POU domain-containing octamer-binding protein | 1,26 | 1,21E-03 |
| P30837 | AL1B1 | ALDH1B1 | Aldehyde dehydrogenase X | 1,98 | 1,50E-03 |
| E9PLA9 | CAPRIN1 | CAPRIN | Caprin-1 | 1,47 | 1,61E-03 |
| P17987 | TCPA | TCP1 | T-complex protein 1 subunit alpha | 1,91 | 2,84E-03 |
| P39019 | RS19 | RPS19 | 40S ribosomal protein S19 | 1,44 | 3,74E-03 |
| O43143 | DHX15 | DHX15 | Putative pre-mRNA-splicing factor ATP-dependent RNA helicase | 1,00 | 4,00E-03 |
| P61586 | RHOA | RHOA | Transforming protein RhoA | 1,01 | 4,49E-03 |
| C9J9K3 | RPSA | RPSA | 40S ribosomal protein SA | 1,07 | 4,91E-03 |
| A0A087WXM6 | RPL17 | RPL17 | 60S ribosomal protein L17 | 1,01 | 5,48E-03 |
| P38919 | EIF4A3 | EIF4A3 | Eukaryotic initiation factor 4A-III | 1,08 | 7,23E-03 |
| P00505 | AATM | GOT2 | Aspartate aminotransferase, mitochondrial | 1,11 | 8,63E-03 |
| P27105 | STOM | STOM | Erythrocyte band 7 integral membrane protein | 1,01 | 9,61E-03 |
| A0A096LNZ9 | ISG15 | ISG15 | Ubiquitin-like protein ISG15 | 1,25 | 1,10E-02 |
| E9PEX6 | DLD | DLD | Dihydrolipoyl dehydrogenase, mitochondrial | 1,08 | 1,66E-02 |
| Q10589-2 | BST2 | BST2 | Bone marrow stromal antigen 2 | 1,15 | 2,37E-02 |
| Q06210 | GFPT1 | GFPT1 | Glutamine--fructose-6-phosphate aminotransferase | 3,22 | 2,49E-02 |
| A0A087WVM3 | CYR61 | CYR61 | Protein CYR61 | 1,34 | 2,75E-02 |
| Q13162 | PRDX4 | PRDX4 | Peroxiredoxin-4 | 1,13 | 2,88E-02 |

**Table S2: Primer sequences used for siRNA, qPCR and mutagenesis**

| Experiment | Target | Sequence |
|---|---|---|
| siRNA | TG2 | GGGCGAACCACCUGAACAAdTdT |
| | GFPT1.1 | GACAGAUUGUGGAGUUCAUdTdT |
| | GFPT1.2 | CCUUGGUGGAGAGAGUUAUdTdT |
| | GFPT1.3 | GUGACUUCCUGGACAGAAAdTdT |
| | GFPT2.1 | GUUCCAAGUUUGCGUAUAAdTdT |
| | GFPT2.2 | GACCGAAUUUCACUACAAAdTdT |
| | GFPT2.3 | CCAUCGCCAAGCUGAUUAAdTdT |
| qPCR | α-actin | GGACTTCGAGCAAGAGATGG |
| | | GCAGTGATCTCCTTCTGCATC |
| | *Chlamydia* 16S RNA | TGGATGAGGCATGCAAGTA |
| | | TACTAACCCTTCCGCCACTAAA |
| | Mouse 18S RNA | TAACGAACGAGACTCTGGCAT |
| | | CGGACATCTAAGGGCATCACAG |
| | TG2 | TAAGAGATGCTGTGGAGGAG |
| | | CGAGCCCTGGTAGATAAA |
| | GLUT-1 | AACTCTTCAGCCAGGGTCCAC |
| | | CACAGTGAAGATGATGAAGAC |
| | GLUT-3 | CACAGGTTTTGTGCCCATGTA |
| | | ACCCCGCAGGGCAGTAG |
| Mutagenesis | Q58N | GGGAAGCCAATGCCTGCAAAATCAATCTTATTAAGAAGAAAGGAAAAGT |
| | | ACTTTTCCTTTCTTCTTAATAAGTTGATTTTGCAGGCATTGGCTTCCC |
| | Q326N | CGAGCTGTGCAAACACTCAATATGGAACTCCAGCAGATC |
| | | GATCTGCTGGAGTTCCATATTGAGTGTTTGCACAGCTCG |
| | Q548N | CGAAATTCAGAAACTAGCAACAGAACTTTATCATAATAAGTCAGTTCTGATAATG |
| | | CATTATCAGAACTGACTTATTATGATAAAGTTCTGTTGCTAGTTTCTGAATTTCG |
